# Neuronal calcium sensors emerge as novel adenylyl cyclase 8 regulators

**DOI:** 10.64898/2026.04.19.719435

**Authors:** Basavraj Khanppnavar, Roberta Florea, Dina Schuster, Pia Lavriha, Angela Kosturanova, Marco M. Ruckstuhl, Ilayda Kantarci, Anand Vaithia, Jiuhan Shi, Paola Picotti, Alvar D. Gossert, Alexander Leitner, Volodymyr M. Korkhov

## Abstract

Adenylyl cyclase 8 (AC8) is a Ca^2+^-sensitive membrane adenylyl cyclase highly expressed in the central nervous system. AC8-mediated intracellular cyclic adenosine monophosphate (cAMP) accumulation shapes synaptic function, plasticity, and memory formation and is tightly controlled by intracellular Ca^2+^ and G protein–coupled receptor signaling. Although it is well known that the effects of Ca^2+^ on AC8 activity are directly mediated by calmodulin, it has remained unclear until now whether other Ca^2+^-binding proteins also regulate AC8 function. Here, we identify the neuronal calcium sensor hippocalcin-like protein 1 (HPCAL1) as a direct interaction partner of AC8. Using biochemistry, structural proteomics, integrative modeling, cryo-EM and cell-based FRET approaches, we show that HPCAL1 associates with AC8 in a Ca^2+^-dependent manner. HPCAL1 shows weak positive modulation of AC8 activity *in vitro*, with sub micromolar affinity. HPCAL1 interacts with flexible regulatory regions of AC8, including the C1b domain and the N- and C-terminal regions. Interestingly, other members of the neuronal calcium sensor family can bind AC8 via the same sites. Together, our study reveals a previously unrecognized mode of Ca^2+^-dependent AC8 recognition by a group of neuronal calcium sensor proteins, which may be relevant to context-dependent regulation of cAMP signaling.

## Introduction

Adenylyl cyclases (ACs) catalyze the conversion of adenosine triphosphate (ATP) into cyclic adenosine monophosphate (cAMP), a ubiquitous intracellular second messenger that regulates numerous fundamental cellular processes. Calcium-sensitive membrane ACs are key downstream effectors of G protein–coupled receptor (GPCR) and Ca^2+^ signaling pathways [1–4]. Among these enzymes, membrane AC isoform 8 (AC8) is one of the principal Ca^2+^-sensitive ACs expressed in the mammalian brain, with high abundance in the piriform cortex, hippocampus, dentate gyrus, thalamus, hypothalamus, cerebellum, and olfactory bulb, predominantly in neurons and astrocytes [5]. AC8-mediated cAMP production is tightly regulated by GPCR signaling through heterotrimeric G proteins and by Ca^2+^ signaling via the Ca^2+^-binding protein calmodulin (CaM). By acting as a coincidence detector for hormonal and neurotransmitter inputs together with intracellular Ca^2+^ signals, AC8 enables context-dependent cellular responses that underlie complex physiological processes, including olfactory processing, learning, memory formation, and motor control [1].

Neuronal cAMP and Ca^2+^ signaling pathways are tightly regulated in both space and time, enabling precise and highly dynamic control of downstream signaling events that govern synaptic transmission and plasticity [2]. Cytosolic Ca^2+^ concentrations in neurons are maintained at resting levels of approximately 50–100 nM but can transiently rise into the micromolar range (up to ∼10 μM) during electrical activity [3]. The effective free Ca^2+^ concentration is shaped by Ca^2+^-buffering proteins such as calretinin [4], regulated Ca^2+^ influx through voltage-gated Ca^2+^ channels [5], and Ca^2+^ extrusion mediated by the sodium–calcium exchanger [6]. These dynamic Ca^2+^ fluctuations are decoded by specialized Ca^2+^ sensor proteins, most prominently calmodulin (CaM), which regulates the activity of a broad range of molecular targets, including ion channels and signaling enzymes [7-9]. Notably, CaM frequently pre-associates with its binding partners, including AC8, thereby enabling rapid and graded responses to subtle Ca^2+^ elevations and enhancing the temporal and spatial precision of downstream signaling.

In addition to CaM, neurons express a diverse array of other Ca^2+^-binding proteins that contribute to the specificity, complexity, and compartmentalization of Ca^2+^-dependent signaling [10]. Among these, the neuronal calcium sensor (NCS) protein family represents a major class of Ca^2+^-binding proteins involved in neuronal signaling [11]. The NCS family comprises 14 members, including NCS1, visinin-like proteins (VILIP1–3), recoverin, hippocalcin, neurocalcin-δ, guanylyl cyclase–activating proteins (GCAP1–3), and voltage-gated potassium channel–interacting proteins (KChIP1–4) [12]. Most NCS proteins harbor an N-terminal myristoylation, whereas some members (e.g., KChIP2 and KChIP3) are palmitoylated [15]. Binding of Ca^2+^ to the conserved EF-hand motifs of NCS proteins induces conformational rearrangements that expose their lipid modifications, enabling reversible association with cellular membranes via a Ca^2+^-dependent “myristoyl switch” [17]. This mechanism enables NCS proteins to dynamically engage with membrane-associated signaling complexes. However, not all NCS proteins follow this paradigm. For example, NCS1 and KChIP1 remain membrane-associated even at low Ca^2+^ concentrations [12, 13]. Differences in Ca^2+^ affinity (approximately 200 nM–10 μM), membrane targeting, and conformational dynamics enable NCS proteins to selectively regulate distinct molecular targets and signaling pathways [14].

Previous studies have indicated that there are functional links between neuronal calcium sensors and AC signaling. For example, visinin-like protein 1 (VILIP1) has been reported to modulate AC activity in heterologous expression systems [15] and to inhibit olfactory AC3 in a Ca^2+^-dependent manner following olfactory stimulation [16]. In addition, both myristoylated and myristoylation-deficient forms of VILIP1 have been shown to influence intracellular cAMP levels in C6 glioma cells [17, 18].

Interestingly, GCAPs, a subfamily of neuronal calcium sensor (NCS) proteins are well-established direct regulators of retinal guanylyl cyclases (GCs). In photoreceptors, GCAP1 and GCAP2 bind directly to retinal guanylyl cyclases such as RetGC1, regulating their catalytic activity in a Ca^2+^-dependent manner. Under low Ca^2+^ conditions, GCAPs stimulate cyclase activity; however, Ca^2+^ binding converts them into inhibitory regulators. Biochemical and structural studies indicate that GCAP1 primarily interacts with the intracellular regulatory regions of RetGC1, engaging elements of the kinase homology domain (KHD) and adjacent segments near the dimerization and catalytic domains that transmit allosteric control to the catalytic center. This provides a clear evolutionary precedent for direct enzymatic control of cyclic nucleotide signaling by NCS proteins [19] **(Figure 1A)**. Nevertheless, it remains unclear whether the reported effects of NCS proteins on cAMP signaling are due to direct physical interactions with AC or indirect modulation of upstream GPCR-cAMP signaling components.

**Figure 1.**
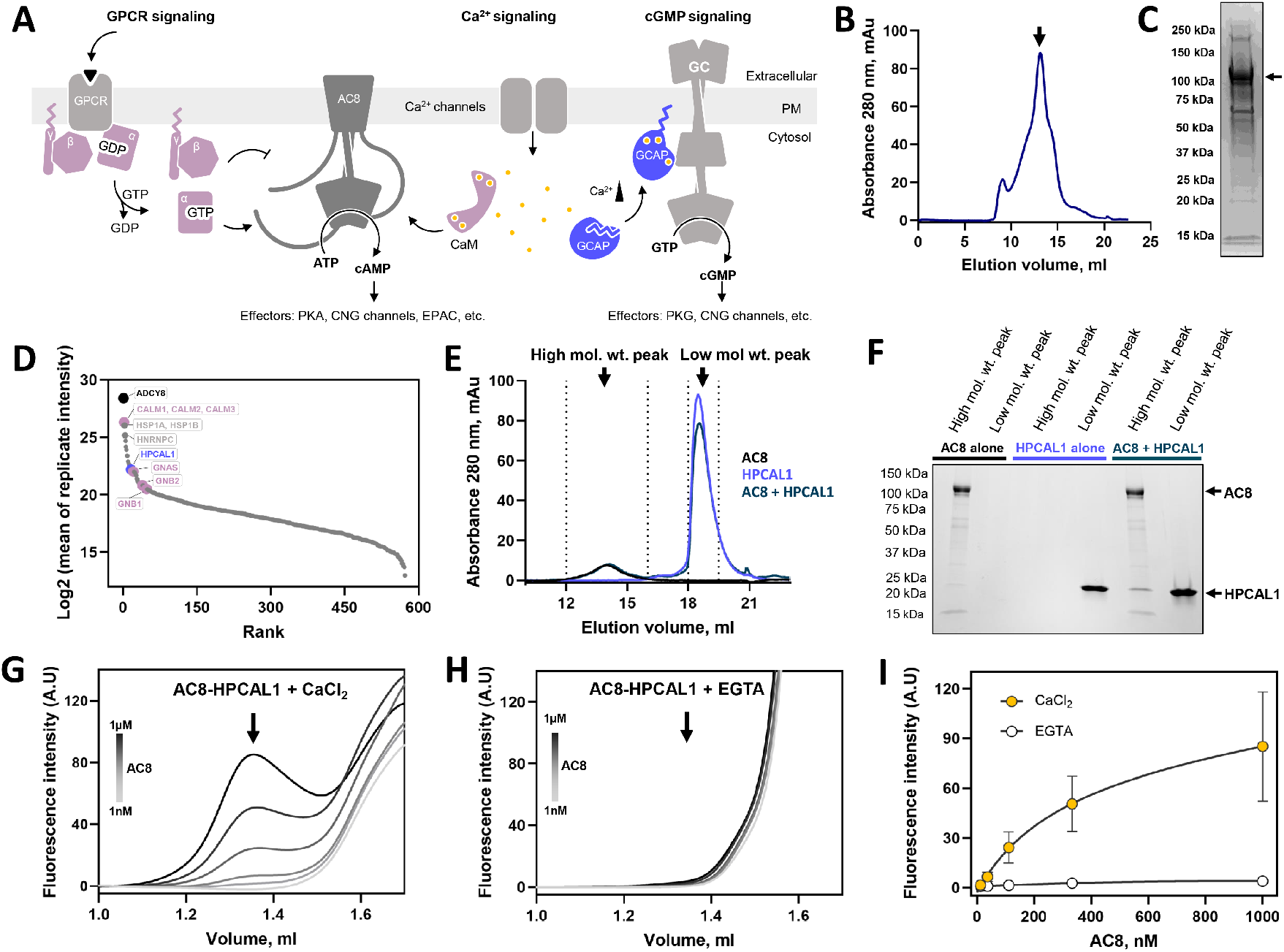
Identification of HPCAL1 as a Ca^2+^-dependent interaction partner of AC8. **(A)** Schematic overview of Ca^2+^- and GPCR-dependent cyclic nucleotide signaling pathways. Activation of G protein–coupled receptors (GPCRs) stimulates AC8 via heterotrimeric G proteins, while Ca^2+^ influx activates calmodulin (CaM) to regulate AC8 activity and cAMP production. In parallel, membrane guanylyl cyclases (GCs) are directly regulated by guanylyl cyclase–activating proteins (GCAPs) in a Ca^2+^-dependent manner, highlighting a conceptual framework for Ca^2+^ sensor–mediated regulation of cyclic nucleotide synthesis. **(B)** Size-exclusion chromatography (SEC) profiles and **(C)** Coomassie-stained SDS–PAGE of affinity-purified recombinant AC8 used for bottom-up proteomics analysis. The arrow indicates the position of full-length AC8. **(D)** Rank-ordered plot of proteins co-purifying with AC8 identified by label-free quantitative mass spectrometry. The known AC8 interactors, including calmodulin and heteromeric G proteins Gαs, are highlighted. HPCAL1 is among the most abundant co-purified proteins, indicating a potential interaction with AC8. **(E)** SEC-based pulldown experiments profiles of pre-incubated AC8–HPCAL1 mixture (dark navy blue), AC8 alone (black), and HPCAL1 alone (blue). Co-elution of HPCAL1 with AC8 in fraction 1 indicates complex formation. **(F)** SDS–PAGE analysis of SEC fractions corresponding to panel (E). HPCAL1 is detected in AC8-containing fractions only when both proteins were pre-incubated, confirming co-elution and supporting direct interaction *in vitro*.

In this study, we identify hippocalcin-like protein 1 (HPCAL1, also known as VILIP3) and other NCS homologs as direct interaction partners of AC8. NCS proteins are expressed in multiple brain regions, including the cerebellum, hippocampus, dentate gyrus, and hippocampal neurons [19], substantially overlapping with regions of high AC8 expression and supporting a role for NCS in the modulation of AC8-dependent signaling. We use integrated structural and functional approaches to elucidate the molecular basis of the Ca^2+^-dependent interaction between AC8 and NCS proteins. Together, our findings reveal NCS proteins as previously unrecognized regulators of AC8 and highlight a new level of Ca^2+^-dependent control in neuronal cAMP signaling.

## Results

### Neuronal calcium sensor HPCAL1 co-purifies with membrane adenylyl cyclase AC8

Building on our previous observation that well-established interactors (e.g. calmodulin and heterotrimeric G proteins) co-purify with recombinant AC8, we performed a standard bottom-up proteomics workflow to identify new AC8 interacting partners. Purified AC8 was subjected to tryptic digestion and analyzed via liquid chromatography-tandem mass spectrometry (LC-MS/MS), which confirmed the presence of known partners alongside molecular chaperones (e.g., Hsp70) and transport proteins such as syntaxin-7 (**Figure 1B-D**). In addition, multiple G protein subunits were detected, including α subunits (Gαs, GNAS; Gα13, GNA13; Gαi2, GNAI2), β subunits (GNB1, GNB2, GNB4), and γ subunits (GNG2, GNG5, GNG12). Having successfully identified established interaction partners in the tryptic digest, we proceeded to screen the dataset for novel regulatory candidates. When proteins were ranked according to their label-free quantification intensities, the neuronal calcium sensor hippocalcin-like protein-1 (HPCAL1) emerged as a highly abundant protein co-purified with AC8 (**Figure 1D**).

The presence of a protein in an affinity-purified sample does not necessarily indicate a direct interaction, as co-purified proteins may arise from non-specific binding to affinity reagents, solid supports, or contamination from other purification experiments [20]. However, the prominent detection of HPCAL1 warranted further investigation, given that HPCAL1 is an NCS protein that is homologous to GCAP1. GCAP1 is a well-established direct regulator of the membrane protein RetGC1. Further, ACs and GCs themselves share partial structural homology (**Figure 1A**). Functional interactions between NCS proteins and ACs have been suggested in previous studies [15-19]. These observations prompted us to perform further experiments to validate HPCAL1 as a bona fide binding partner of AC8.

To directly observe the AC8-HPCAL1 interaction we used size-exclusion chromatography (SEC, **Figure 1E-F**). In these experiments, equal amounts of purified HPCAL1 alone, AC8 alone, or a pre-incubated mixture of the two were injected onto a SEC column. The higher molecular weight elution fractions corresponding to the AC8 peak were collected, concentrated, and analyzed by sodium dodecyl sulfate–polyacrylamide gel electrophoresis (SDS– PAGE, **Figure 1F**). In the presence of HPCAL1, an additional band corresponding to the molecular weight of HPCAL1 was detected in the AC8-containing elution fraction. The absence of this band in the high molecular weight SEC fractions obtained using HPCAL1 alone confirms the formation of an AC8–HPCAL1 complex *in vitro*.

### HPCAL1 binds AC8 in a calcium-dependent manner

The interaction of calcium sensor proteins with their targets is tightly regulated by intracellular Ca^2+^ levels [10]. To test whether HPCAL1 interacts with AC8 in a Ca^2+^-dependent manner, we performed fluorescence-detection size-exclusion chromatography (FSEC) binding assays that we previously used to characterize AC8–calmodulin interactions [21]. Briefly, purified AC8 and a C-terminally cyan fluorescent protein–tagged HPCAL1 (HPCAL1– CFP) were analyzed by SEC. Upon addition of AC8, we observed a concentration-dependent increase in CFP fluorescence signal co-eluting with AC8 at its characteristic elution volume, indicating association of HPCAL1 with AC8, with an apparent dissociation constant (K_d_) of ∼290 nM **(Figure 2A-C)**. In the presence of the Ca^2+^ chelator EGTA, the CFP fluorescence signal at the AC8 elution volume was abolished, even at higher concentrations of AC8, indicating that the interaction between AC8 and HPCAL1 is Ca^2+^ dependent.

**Figure 2.**
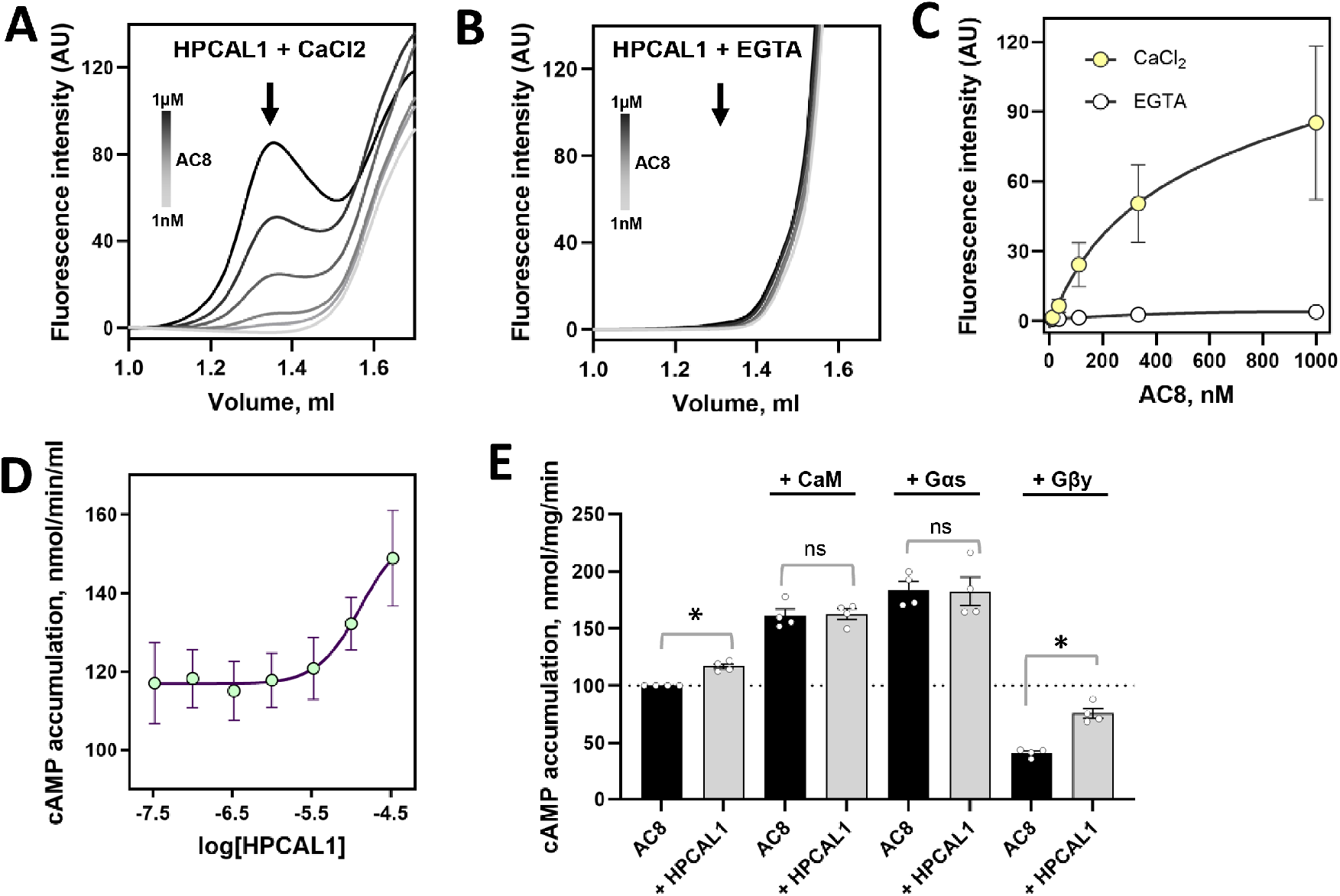
HPCAL1 weakly regulates AC8 activity in Ca^2+-^dependent manner. **(A-B)** Fluorescence-detection SEC (FSEC) analysis of purified AC8 incubated with C-terminally CFP-tagged HPCAL1 (HPCAL1–CFP) in the presence of 0.5 mM CaCl_2_ or 0.5 mM Ca^2+^ chelator EGTA. **(C)** Quantification of CFP fluorescence intensity at the AC8 elution peak as a function of AC8 concentration in the presence of Ca^2+^ (purple) or EGTA (gray), yielding an apparent dissociation constant (*Kd*) of ∼292 nM under Ca^2+^-replete conditions. Data shown as mean ± SEM obtained from 5 independent experimental replicates. **(D)** *In vitro* cAMP accumulation assay showing the effect of increasing concentrations of HPCAL1 on AC8 activity in the presence of Ca^2+^. **(E)** The impact of HPCAL1 on regulation of AC8 by G protein βγ subunits (Gβγ), G protein α subunit (Gαs), and calmodulin (CaM). HPCAL1 alone weakly modulates AC8 activity and does not measurably alter CaM-dependent activation under these assay conditions. HPCAL1 does not significantly affect CaM- or Gαs-mediated stimulation of AC8 but could affect Gβγ-dependent inhibition of AC8. Data for cAMP assays are shown as mean ± SEM obtained from 4 independent experimental replicates. Statistical significance is indicated by asterisks (*) with p value < 0.01, while ‘ns’ denotes no significant difference, as determined by one-way ANOVA.

Next, we tested whether HPCAL1 could modulate AC8 activity, analogous to the regulation of GCs by GCAP1. The *in vitro* cAMP accumulation assays performed in the presence of 0.5 mM CaCl_2_ revealed weak activation of AC8 by HPCAL1 (**Figure 2D**). As HPCAL1 did not substantially modulate AC8 activity, we next tested whether it’s binding competitively affects the regulation of AC8 by other established regulatory partners. Our results show that HPCAL1 can partially relieve Gβγ-mediated inhibition of AC8, whereas it does not measurably influence CaM- or Gαs-mediated stimulation of AC8 (**Figure 2E**). These observations suggest that HPCAL1 may act as a context-dependent modulator of AC8 signaling, particularly in compartments where strong activators such as CaM or Gαs are absent.

### Cryo-EM analysis indicates that HPCAL1 interacts with flexible regions of AC8

To structurally characterize the interaction between AC8 and HPCAL1, we performed single-particle cryo-electron microscopy (cryo-EM) analysis of the complex in the presence and absence of Gαs. Cryo-EM datasets were collected for four conditions: AC8 alone, AC8–HPCAL1, AC8–Gαs, and the ternary AC8–Gαs–HPCAL1 complex (Figure 3). The concentrations of each of the protein components (AC8, HPCAL1, Gαs) in the cryo-EM samples were controlled by mixing the purified proteins at specific molar ratios prior to grid freezing. This included a 2.5-fold molar excess of HPCAL1 and a 1.2-fold molar excess of Gαs relative to AC8. All four datasets were prepared from the same batch of purified proteins, and comparable amounts of cryo-EM data (∼ 3000 movies each) were collected and processed using identical workflows to enable direct comparison between conditions (**Supplementary Figures S1)**.

**Figure 3.**
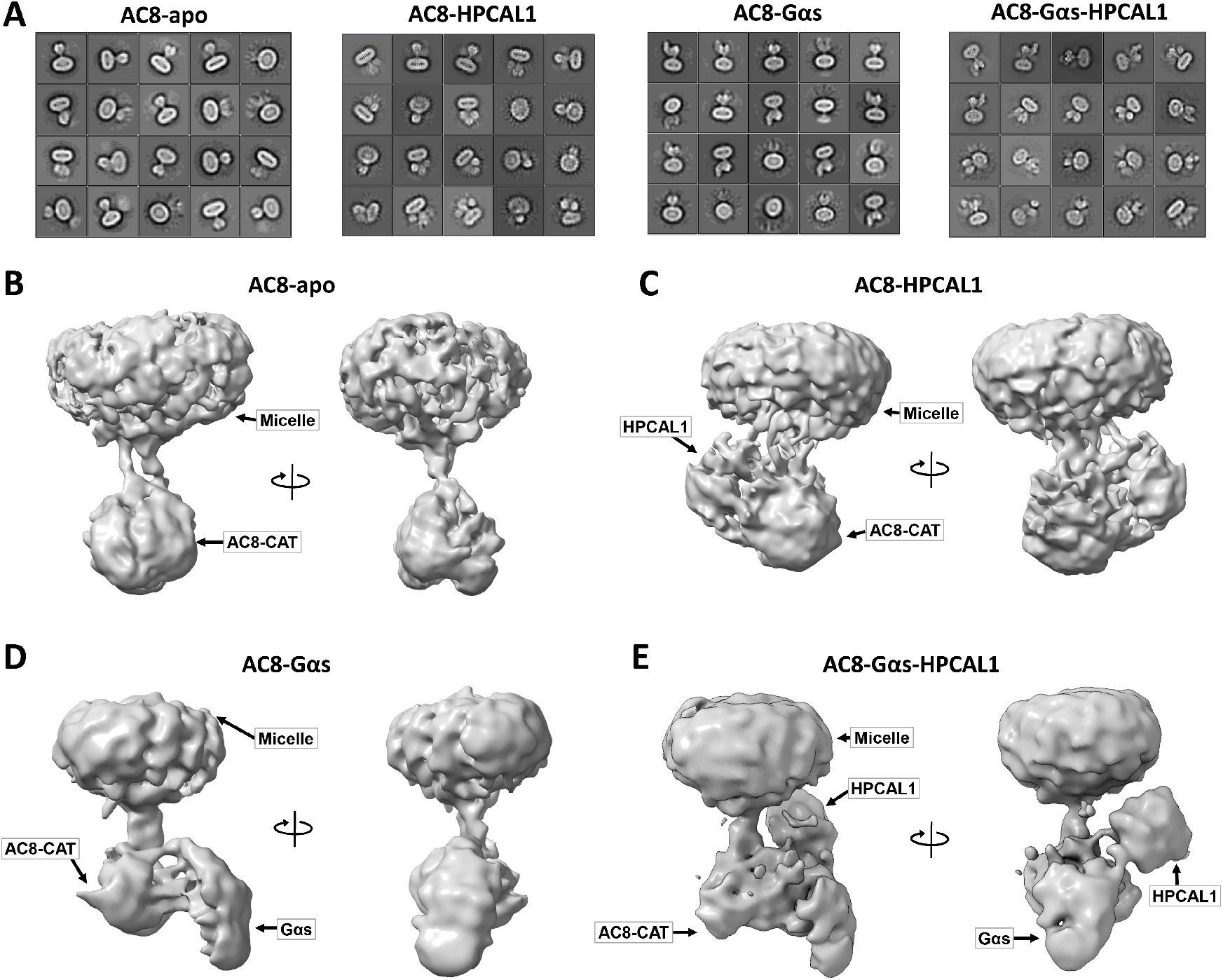
Cryo-EM analysis of AC8-HPCAL1 complexes. **(A)** Representative 2D class averages of AC8 in in presence and absence of Gαs and HPCAL1. **(B-E)** Low-resolution cryo-EM maps of the AC8 alone **(B)**, AC8– HPCAL1 complex **(C)**, AC8–Gαs **(D)** and the AC8–Gαs–HPCAL1 complex **(E)**. In the presence of HPCAL1, an additional density is observed above the AC8 catalytic domain (CAT) and adjacent to the helical domain.

Two-dimensional (2D) class averages revealed clear differences between the datasets. In particular, the AC8– HPCAL1 and AC8–Gαs–HPCAL1 complexes displayed additional globular density compared with the AC8-apo and AC8–Gαs datasets (**Figure 3A**), suggesting the association of HPCAL1. Three-dimensional (3D) reconstructions further supported this interpretation. In the AC8-apo reconstruction, only the micelle-associated transmembrane region and the catalytic domain are resolved (**Figure 3B**). Addition of HPCAL1 gives rise to extra density adjacent to the cytoplasmic region connecting the transmembrane and catalytic domains (**Figure 3C**). In the presence of Gαs, a distinct density corresponding to Gαs is observed engaging the catalytic domain of AC8 (**Figure 3D**). When both regulators are present, densities attributable to both Gαs and HPCAL1 are observed near the cytoplasmic region of AC8 (**Figure 3E**).

Based on these comparative observations, larger cryo-EM datasets were subsequently collected for the AC8– HPCAL1 and AC8–Gαs–HPCAL1 complexes and processed extensively to obtain high-resolution cryo-EM reconstructions. Three additional datasets comprising 8,974, 13,588, and 11,360 movies were acquired for the AC8–HPCAL1 complex, (**Supplementary Figures S2)**. and an additional dataset of 6,363 movies was collected for the AC8–Gαs–HPCAL1 complex (**Supplementary Figures S3)**. Despite these extensive efforts, including multiple rounds of 3D classification, determining a high-resolution structure of AC8-bound HPCAL1 remained unsuccessful. This limitation likely reflects pronounced conformational flexibility of the cytoplasmic regions of AC8 involved in HPCAL1 binding.

### Full-length AC8 is important for interaction with HPCAL1

As high-resolution cryo-EM structure determination of the AC8–HPCAL1 complex was unsuccessful, we next sought to delineate the AC8 regions required for interaction with HPCAL1 using a domain-deletion strategy. AC8 constructs lacking the N-terminus (Nt), C-terminus (Ct), or C1b domain were generated and analysed. However, these AC8 deletion constructs were highly unstable and could not be expressed or purified robustly **(Supplementary Fig. S4)**.

As an alternative strategy, the AC8 Nt, Ct, and C1b peptides or domains were expressed and purified individually **(Supplementary Fig. S5)**, after which their interactions with HPCAL1 were analyzed using nuclear magnetic resonance (NMR) spectroscopy. Under these conditions, no detectable interactions between HPCAL1 and the isolated AC8 Nt, Ct, or C1b regions were observed, judged by 1H–15N HSQC spectra **(Supplementary Fig. S6)**. These results suggest that the interaction between AC8 and HPCAL1 requires a specific three-dimensional arrangement of the protein domains in the context of a full-length AC8 that is not recapitulated by isolated domains or peptides.

### Interaction between AC8 and HPCAL1 in cell-based assays

Encouraged by our *in vitro* findings on purified AC8 and HPCAL1, we decided to assess whether these proteins interact in a cellular environment. As a prerequisite, we first tested whether HPCAL1, similar to other NCS proteins, undergoes a Ca^2+^-myristoyl switch that promotes its translocation from the cytosol to the plasma membrane upon elevation of intracellular Ca^2+^ levels[12]. To assess whether Ca^2+^-dependent plasma membrane recruitment of HPCAL1 could be induced and detected in mammalian cells, CFP-tagged HPCAL1 was expressed in HEK293F cells and cells were treated with ionomycin, a Ca^2+^ ionophore that increases intracellular Ca^2+^ concentrations by releasing Ca^2+^ from intracellular stores and activating store-operated Ca^2+^ entry[22, 23].

Time-resolved imaging revealed a redistribution of HPCAL1 from the cytosol to the plasma membrane following ionomycin addition **(Figure 4A)**. To confirm that ionomycin treatment elevated intracellular Ca^2+^ levels under these conditions, cells expressing the Ca^2+^ sensor G-GECO were analyzed in parallel. Ionomycin treatment resulted in a robust increase in G-GECO fluorescence, indicating elevated intracellular Ca^2+^ levels **(Figure 4B)**. Quantification of HPCAL1 fluorescence under the same conditions showed a decrease in cytosolic signal accompanied by an increase in membrane-associated fluorescence over time (Figure 4C), consistent with activation of the Ca^2+^-myristoyl switch **(Figure 4C)**.

**Figure 4.**
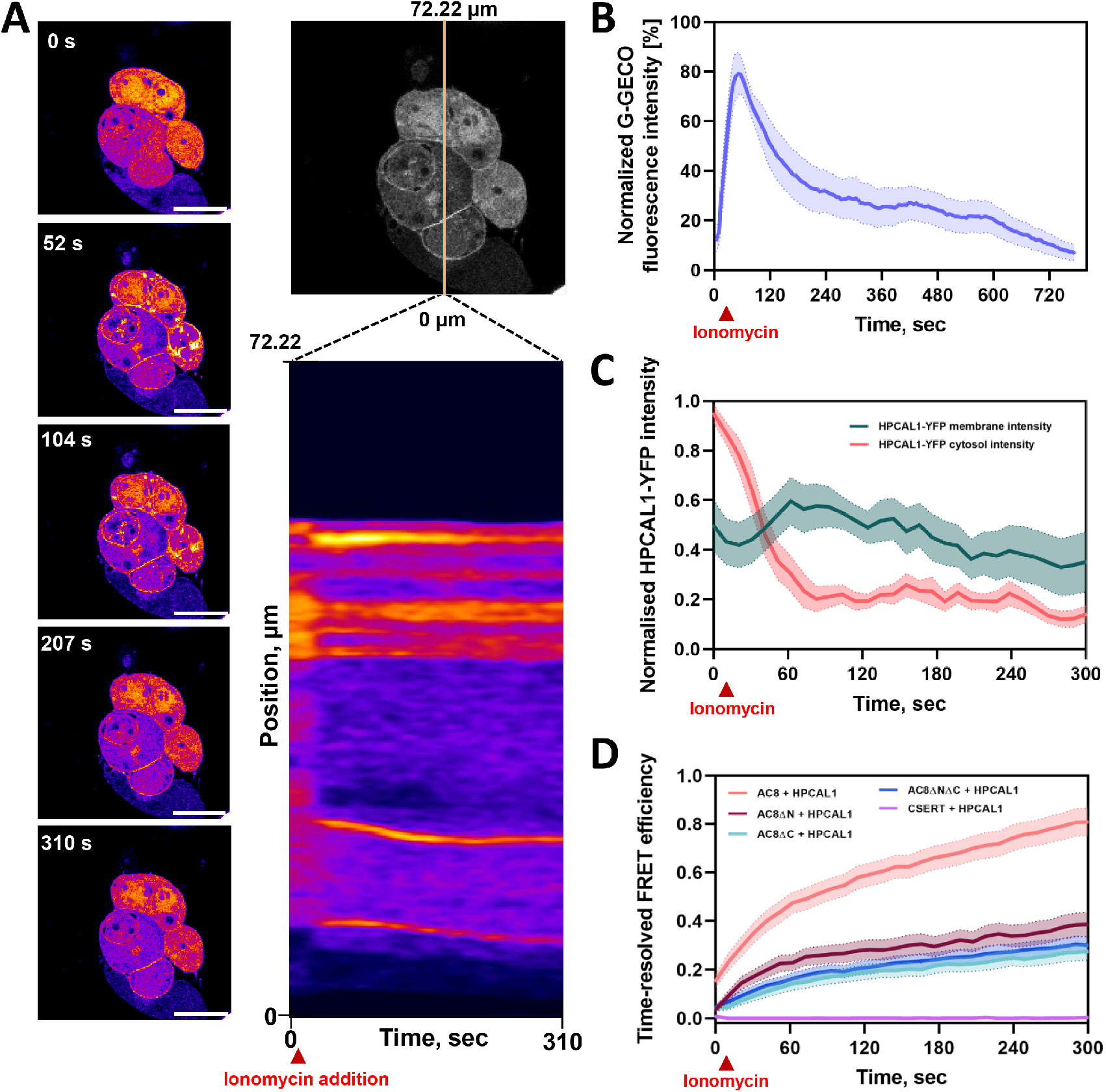
Synergistic roles of AC8 flexible domains for interaction with HPCAL1. **(A)** Representative time-resolved fluorescence images of a HPCAL1–YFP transfected HEK293F cell following ionomycin treatment (3 μM ionomycin in 10 mM CaCl_2_). The images and corresponding line-scan analysis show increasing fluorescence intensity at the plasma membrane and a decrease in cytosolic fluorescence over time, indicating redistribution of HPCAL1 upon Ca^2+^ elevation. Scale bar, 20 µm. **(B)** Control experiment showing change in intracellular Ca^2+^ levels following ionomycin treatment, measured using the Ca^2+^ sensor G-GECO (mean ± SEM; n = 12). **(C)** Quantification of HPCAL1–YFP fluorescence intensity in the cytosol (orange) and at the plasma membrane (light blue) following ionomycin treatment (mean ± SEM; n_cytosol = 12, n_membrane = 13). **(D)** Time-resolved FRET interaction study of AC8 and HPCAL1 upon calcium stimulation with ionomycin. Wild type AC8 (green) shows the highest FRET efficiency. AC8 truncations (no N-terminus, no C-terminus, no N- and no C-terminus) show reduced FRET efficiency. No FRET signal was detected with the negative control CSERT (CFP-tagged serotonin transporter).

Next, to directly assess whether AC8 and HPCAL1 interact in cells, HEK293F cells were co-transfected with YFP-tagged AC8 and CFP-tagged HPCAL1 and analyzed their interaction using Förster resonance energy transfer (FRET). FRET efficiency was monitored before and after ionomycin-induced Ca^2+^ elevation. Upon ionomycin treatment, a clear increase in FRET between AC8 and HPCAL1 was observed, indicating Ca^2+^-dependent proximity of the two proteins in cells **(Figure 4D)**. As a negative control, the CFP-tagged serotonin transporter (SERT) did not produce a detectable FRET signal with HPCAL1, confirming the specificity of the interaction. The FRET increase was maximal for wild-type AC8 and was reduced when AC8 variants lacking either the N-terminus, the C-terminus, or both were analyzed, although the signal was not completely abolished **(Figure 4D)**.

Thus, although the truncated AC8 variants could not be purified for *in vitro* interaction studies, the cell-based FRET experiments confirm a Ca^2+^-dependent association between AC8 and HPCAL1 in living cells and indicate that both the N-and C-terminal regions of AC8 contribute to this interaction.

### Mass spectrometry-based structural proteomic approaches map the interaction between AC8 and HPCAL1

As the flexibility of the AC8-HPCAL1 complex made it difficult to determine its cryo-EM structure, we performed cross-linking mass spectrometry (XL-MS) to gain further insights into the AC8 regions involved in HPCAL1 binding [24]. Purified AC8 and HPCAL1 were crosslinked using a primary amine-reactive cross-linking reagent, disuccinimidyl suberate (DSS), followed by LC–MS/MS analysis. XL-MS identified intermolecular crosslinks involving both the C1b domain and the N-terminal region of AC8, indicating that HPCAL1 engages multiple flexible segments of AC8 **(Figure 5A)**.

**Figure 5.**
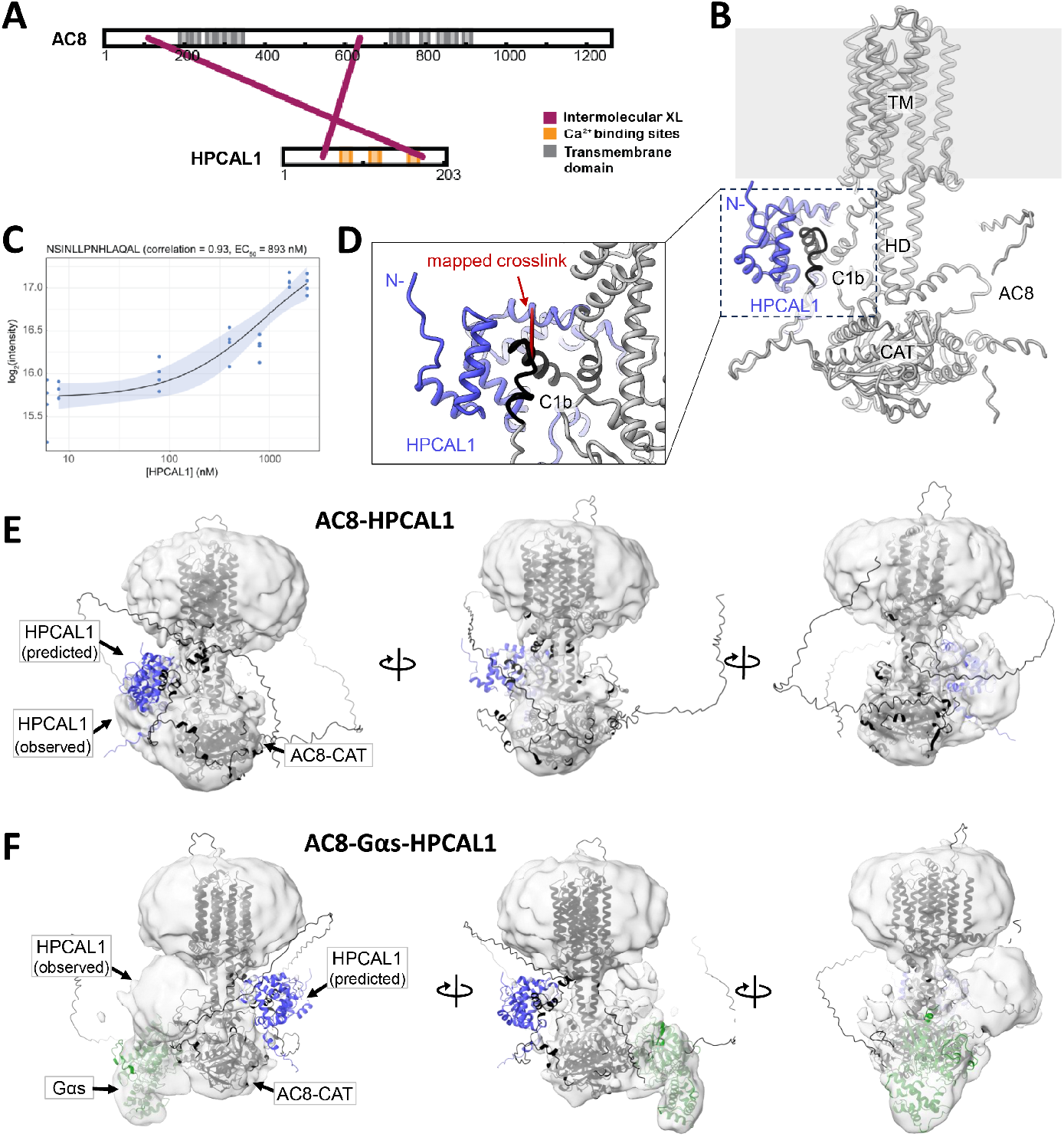
Integrative modeling of the AC8–HPCAL1 interaction. **(A)** XL-MS analysis identifies contacts between HPCAL1 and multiple flexible regions of AC8, including the N-terminal and C1a regions (indicated by purple lines). **(B)** Structural model of the AC8-HPCAL1 complex generated using AlphaLink2. AC8 is shown in gray and HPCAL1 in blue. **(C)** LiP-MS analysis of AC8 in the presence of increasing HPCAL1 concentrations reveals altered protease accessibility of a C1b-derived peptide (NSINLLPNHLAQAL) consistent with **(D) the** predicted model. The apparent interaction affinity is in the submicromolar range, with an EC_50_ of 893 nM and a Pearson correlation coefficient of 0.93 for the dose–response curve. **(E-F)** Overlay of the AlphaLink2 based AC8-HPCAL1 model onto low-resolution cryo-EM reconstructions of the AC8–HPCAL1 complex (panel E) and the AC8–Gαs–HPCAL1 complex (panel F). The AC8–HPCAL1 model could not be unambiguously fitted into the cryo-EM map of the AC8–HPCAL1 complex alone, whereas the presence of Gαs provides additional structural features that allow more reliable placement and interpretation of the model.

To investigate the AC8–HPCAL1 interaction in a native membrane context, we additionally performed limited proteolysis–coupled mass spectrometry (LiP-MS) [25] using membrane suspensions from AC8-overexpressing HEK293 cells, as described previously [21, 26]. Purified HPCAL1 was added to membrane suspensions at increasing concentrations (0–2.4 μM) in the presence of 1 mM CaCl_2_. Samples were subjected to pulse proteolysis using proteinase K, followed by tryptic digestion, and subsequent analysis by LC–MS/MS. Four-parameter dose– response curves were fitted to all quantified peptide intensities [27].

A significant threshold of Pearson correlation coefficient *r* > 0.85 was applied to identify regions exhibiting significant changes in protease accessibility, based on previously established benchmarking studies [26]. We identified a single region within the AC8 C1b domain that exhibited a significant change in protease accessibility upon the addition of HPCAL1. This complements the acquired proximity data from XL-MS. **(Figure 5C)**.

### Integrative modeling reveals mode of AC8-HPCAL1 interaction

To gain structural insight into the AC8–HPCAL1 interaction, we first performed AlphaFold 2- and AlphaFold 3-based complex predictions. However, most of the predicted models had low confidence scores, probably due to a bias against membrane proteins in the AlphaFold training data, as well as an absence of co-evolutionary signals for the specific protein pairs (Table S1). Additionally, the interactions appear to be motif-driven, which has been reported to present challenges in AlphaFold-Multimer predictions [28]

To overcome this limitation, we incorporated experimentally derived XL-MS restraints and performed integrative modeling using AlphaLink2, a method that allows for the incorporation of experimental data to guide structural predictions [29]. While the pLDDT scores remained in the 50–70 range, likely reflecting the inherent flexibility and ‘disorder-to-order’ transition of the AC8 binding motif, the XL-MS-guided models showed substantially improved confidence scores and converged on a consistent interaction mode. (**Figure 5B & D, Supplementary Fig. 7A**). In the highest scoring models, ranked on predicted alignment error (PAE), predicted template modelling (pTM) and interface predicted template modelling (ipTM), HPCAL1 primarily engages the C1b domain of AC8. This is consistent with our LiP-MS data identifying this region as exhibiting altered protease accessibility upon HPCAL1 binding (**Figure 5C**). The predicted orientation positions HPCAL1 above the catalytic core of AC8, with its N-terminal myristoylation oriented toward the membrane-facing side of the complex. This is consistent with the Ca^2+^-dependent membrane recruitment observed in cell-based assays (**Figure 4**).

While the general placement of HPCAL1 relative to AC8 was in line with the low-resolution cryo-EM maps, where HPCAL1 was found near the helical domain and located between the catalytic and transmembrane domain (**Figure 3B-E and 5B**), our predicted AC8-HPCAL1 model could not be clearly fitted into the extra density seen in AC8-HPCAL1 or AC8–HPCAL1–Gαs reconstructions (**Figure 5E-F**). This discrepancy likely reflects substantial conformational heterogeneity of AC8-HPCAL1, rather than an alternative binding mode. Moreover, the predicted models do not involve contact between HPCAL1 and the N- or C-terminal regions of AC8, which were implicated for stable AC8-HPCAL1 interaction by functional and cell-based assays. These flexible regions may therefore contribute transient or dynamic contacts that constrain HPCAL1 positioning in cryo-EM reconstructions but are not captured in static structural models (**Figure 5E-F**). Together, these results support a model in which HPCAL1 engages AC8 through the structured C1b domain while remaining dynamically associated via flexible regions, resulting in a conformationally heterogeneous complex.

### NCS protein family members interact with AC8

Due to the high sequence and structural similarity of NCS proteins, we reasoned that HPCAL1 may not be the only NCS homologue interacting with AC8. To test this, we expressed and purified five other NCS proteins, including HPCAL4, VILIP1, HPCA, NCALD, and NCS1, and assessed their effects on AC8 by performing *in vitro* cAMP accumulation assays. Among the proteins tested, only HPCAL4 elicited weak but significant activation of AC8 that was qualitatively similar to that observed for HPCAL1, whereas no robust activation of AC8 was detected for the other NCS proteins (**Figure 6A**).

**Figure 6.**
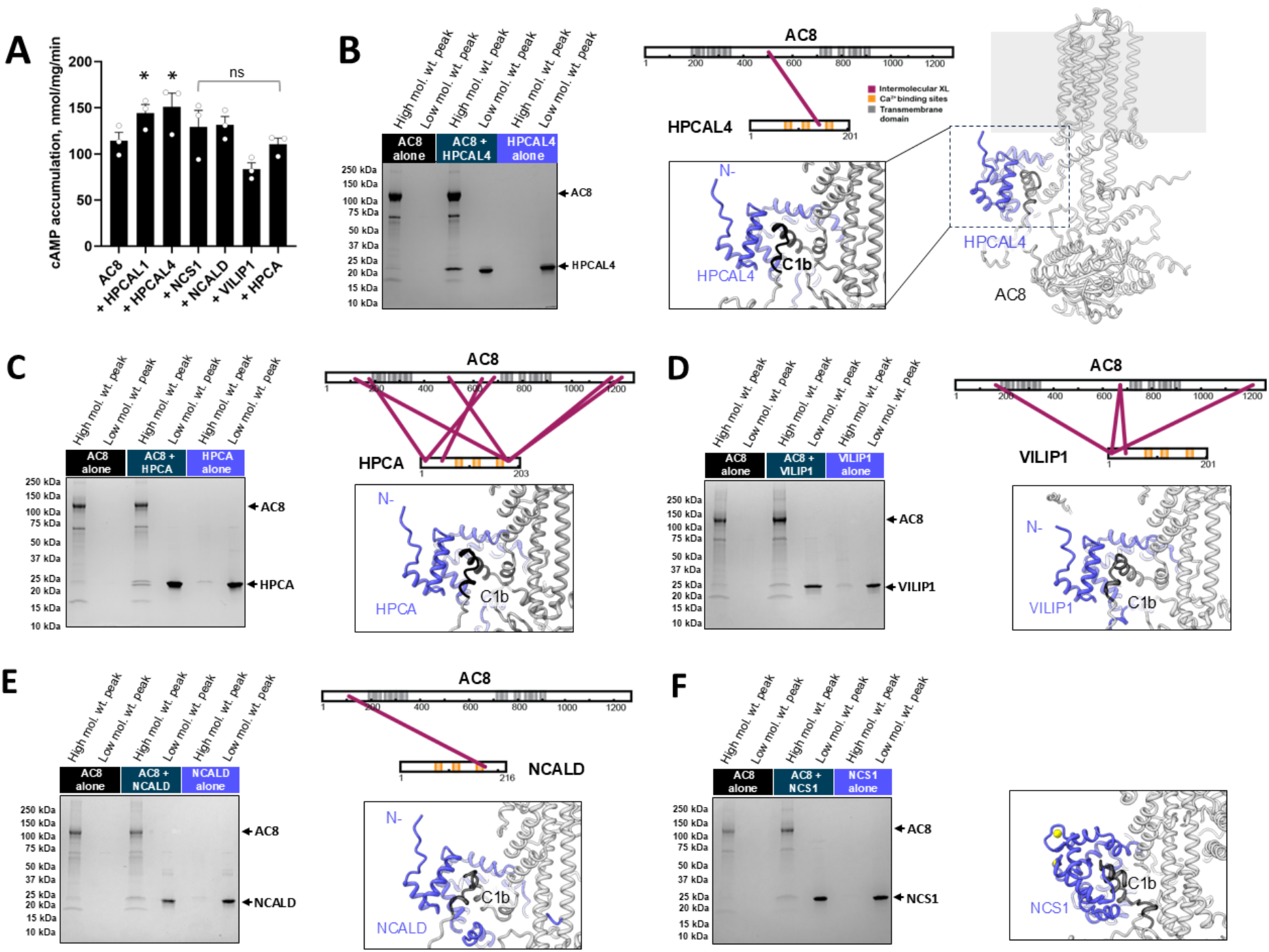
Other members of the neuronal calcium sensor family weakly interact with AC8. **(A)** *In vitro* cAMP accumulation assays measuring AC8 activity in the presence of individual NCS proteins. While most NCS proteins do not significantly alter AC8 activity, HPCAL4 induces a weak activation qualitatively similar to that observed for HPCAL1. Data are shown as mean ± SEM from four independent experiments. Statistical significance is indicated by asterisks (*), whereas “ns” denotes no significant difference relative to basal AC8 activity, as determined by one-way ANOVA. **(B–F)** SEC-based *in vitro* binding assays assessing interactions between AC8 and HPCAL4 (B), HPCA (C), VILIP1 (D), NCALD (E), and NCS1 (F). Binding was evaluated by co-elution of NCS proteins with AC8 in higher–molecular weight fractions compared to the individual proteins alone, following the same analytical strategy described previously for HPCAL1. For each panel, the upper right side depicts experimentally identified crosslinks between AC8 and the respective NCS protein. The lower right corner shows a close-up of the AlphaLink2 models, highlighting the predicted binding site. Crosslinks map predominantly to flexible regions of AC8, including the C1b domain and the N- and C-terminal segments, overlapping with interaction sites identified for HPCAL1. In the case of NCS1, no crosslinks were detected; therefore, the AC8– NCS1 complex was modeled using AlphaFold3. Nevertheless, the predicted models consistently indicate that the hydrophobic binding cleft of NCS proteins engages an amphipathic helix within the AC8 C1b domain.

To determine whether these NCS proteins could nonetheless form stable or transient interactions with AC8, we performed SEC-based *in vitro* binding assays using purified AC8 in combination with each NCS protein. Several of the tested NCS proteins exhibited HPCAL1-like co-elution behavior with AC8, indicative of low-affinity or transient interactions (**Supplementary Fig. S7**). These observations were further supported by XL-MS analysis, which identified crosslinks between AC8 and multiple NCS proteins, except for NCS1 (**Figure 6B-F**).

Notably, the identified crosslinks mapped to similar regions of AC8 as those observed for HPCAL1, primarily involving flexible segments such as the N-terminal region and the C1b domain. In the case of HPCA, additional crosslinks involving the C-terminal region of AC8 were observed. These crosslinks were absent for HPCAL1 but are consistent with cell-based FRET experiments, which indicated an essential contribution of the AC8 C-terminus to the interaction. Using these XL-MS restraints, we performed structure predictions for AC8 bound to other NCS proteins (HPCAL4, HPCA, NCALD and VILIP1) using the AlphaLink2 pipeline. A similar trend of increasing confidence scores was observed, with the AC8 C1b domain occupying the hydrophobic binding cleft of NCS proteins (**Figure 6** and **Supplementary Fig. S8-9, Table S1**).

### Common features in the predicted AC8-NCS complex models

Inspection of the predicted AC8–NCS complex models revealed common structural features across the NCS protein family. In all models, the AC8 C1b segment adopts an amphipathic α-helical conformation positioned within the hydrophobic binding cleft of the NCS proteins (**Figure 7A–B**). The interface is dominated by leucine-rich residues from AC8 that pack against conserved hydrophobic residues lining the NCS cavity.

Structural inspection of the interface further identified serine and threonine residues within the C1b helix positioned at the predicted interaction surface (**Figure 7C**). To evaluate their potential phosphorylation propensity, sequence-based prediction was performed using NetPhos 3.1 [30]. This analysis identified S648 as the most likely phosphorylation site (score 0.807), whereas neighboring threonine residues showed substantially weaker scores (0.43).

Sequence alignment of AC8 orthologs from multiple vertebrate species revealed strong conservation of the C1b segment, including the leucine residues forming the hydrophobic face of the helix as well as the predicted phosphorylation site (**Figure 7D**). Similarly, alignment of NCS proteins highlights conservation of aromatic and hydrophobic residues that form the receptor-binding pocket (**Figure 7E**). Together, these structural and sequence features support a conserved mode of interaction between the AC8 C1b region and the hydrophobic binding cleft of NCS proteins.

**Figure 7.**
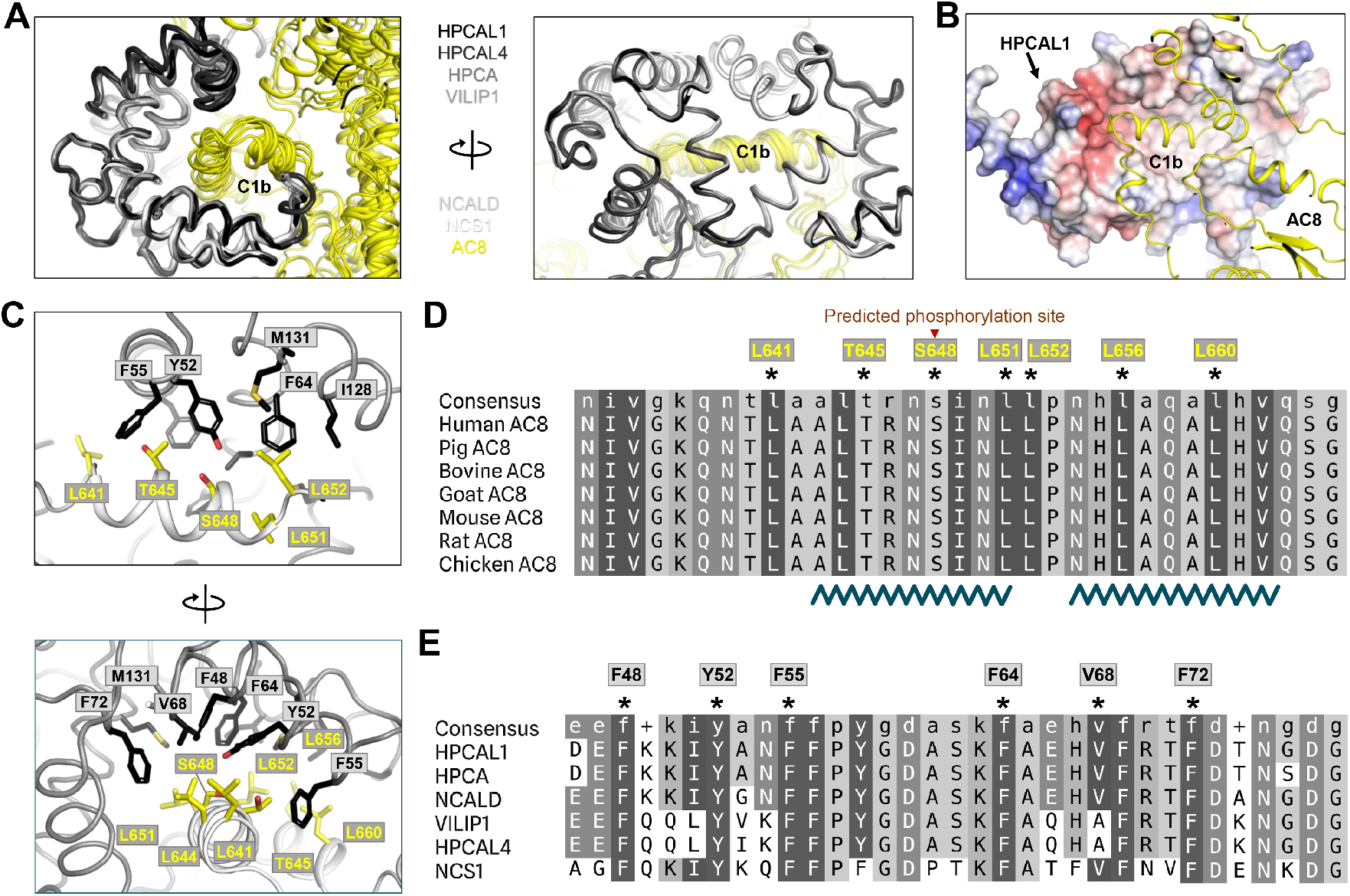
Structural basis of NCS recognition of the AC8. **(A)** Superposition of Alpha-link predicted AC8-NCS complexes highlighting conserved binding mode (HPCAL1, HPCAL4, HPCA, VILIP1, NCALD and NCS1; grey to black, AC8; yellow). **(B)** Electrostatic surface representation of HPCAL1 illustrating its deep hydrophobic binding cleft involved in interaction with AC8 C1b (yellow). **(C)** Close-up view of the AC8–HPCAL1 interface showing key contact residues between AC8 C1b helix (L641, L644, L651, L652, L656, etc.; yellow sticks) and HPCAL1 (e.g., F48, Y52, F55, F64, M131, I128; black sticks). **(D)** Multiple sequence alignment of the AC8 C1b region across vertebrate species (human, pig, bovine, goat, mouse, rat, chicken) demonstrating strong conservation of the leucine-rich amphipathic helices (residue 630-660), and potential phosphorylation site (asterisk), suggesting potential regulation of NCS binding via post-translational modification. **(E)** Sequence alignment of NCS family members (HPCAL1, HPCA, NCALD, VILIP1, HPCAL4, NCS1) highlighting conservation of residues that line the hydrophobic binding groove. Asterisks denote interface residues identified in structural models (F48, Y52, F55, F64, V68, F72), supporting a shared recognition mechanism among NCS proteins.

## Discussion

Protein–protein interactions are fundamental to nearly all aspects of cellular function, including signaling and regulation. Although many interactions have been characterized, transient and low-affinity interactions that underpin dynamic cellular processes have remained understudied due to significant technical challenges in detecting them [31]. In this study, we address these difficulties by combining biochemical assays and cell-based measurements with integrative structural modelling. This multi-pronged approach allowed for the identification and characterisation of NCS proteins as a novel class of transient AC8 interactors.

Our models suggest that NCS proteins recognize a highly conserved, leucine-rich, amphipathic α-helix within the C1b domain of AC8. Although this interaction involves regulatory domains that are specific to AC8, the binding mode is reminiscent of the way in which GCAPs interact with RetGCs. In this case, GCAPs bind to the GC-specific kinase homology domain and to adjacent segments that are located near the dimerization and catalytic domains. These segments transmit allosteric control to the catalytic centre [32-34]. Given that membrane ACs and GCs likely diverged from a common ancestral class III nucleotidyl cyclase [32], this structural similarity hints at a conserved mechanism for calcium sensor-mediated regulation across different cyclases, even if the specific protein domains involved in the process are distinct and the downstream functional outcomes have diverged.

Notably, in contrast to some of the established AC regulators, such as calmodulin and G protein subunits, HPCAL1 and other NCS proteins did not potently modulate AC8 enzymatic activity *in vitro*. Our findings suggest a more nuanced modulatory role. For example, HPCAL1 may influence Gβγ-mediated inhibition of AC, with resulting implications for cAMP signalling in certain cellular contexts. HPCAL1 may counteract the reduction in cAMP levels caused by Gβγ within signalling compartments associated with non-Gαs-coupled receptors (e.g. Gq-coupled receptors). By shielding AC8 from Gβγ, NCS proteins could facilitate crosstalk between GPCR and Ca^2+^ signaling [35]. The predicted overlap between the NCS-binding region and previously proposed Gβγ interaction sites further suggests that these interactions may be mutually exclusive, providing a potential mechanism for regulatory crosstalk (**Figure 8**).

**Figure 8.**
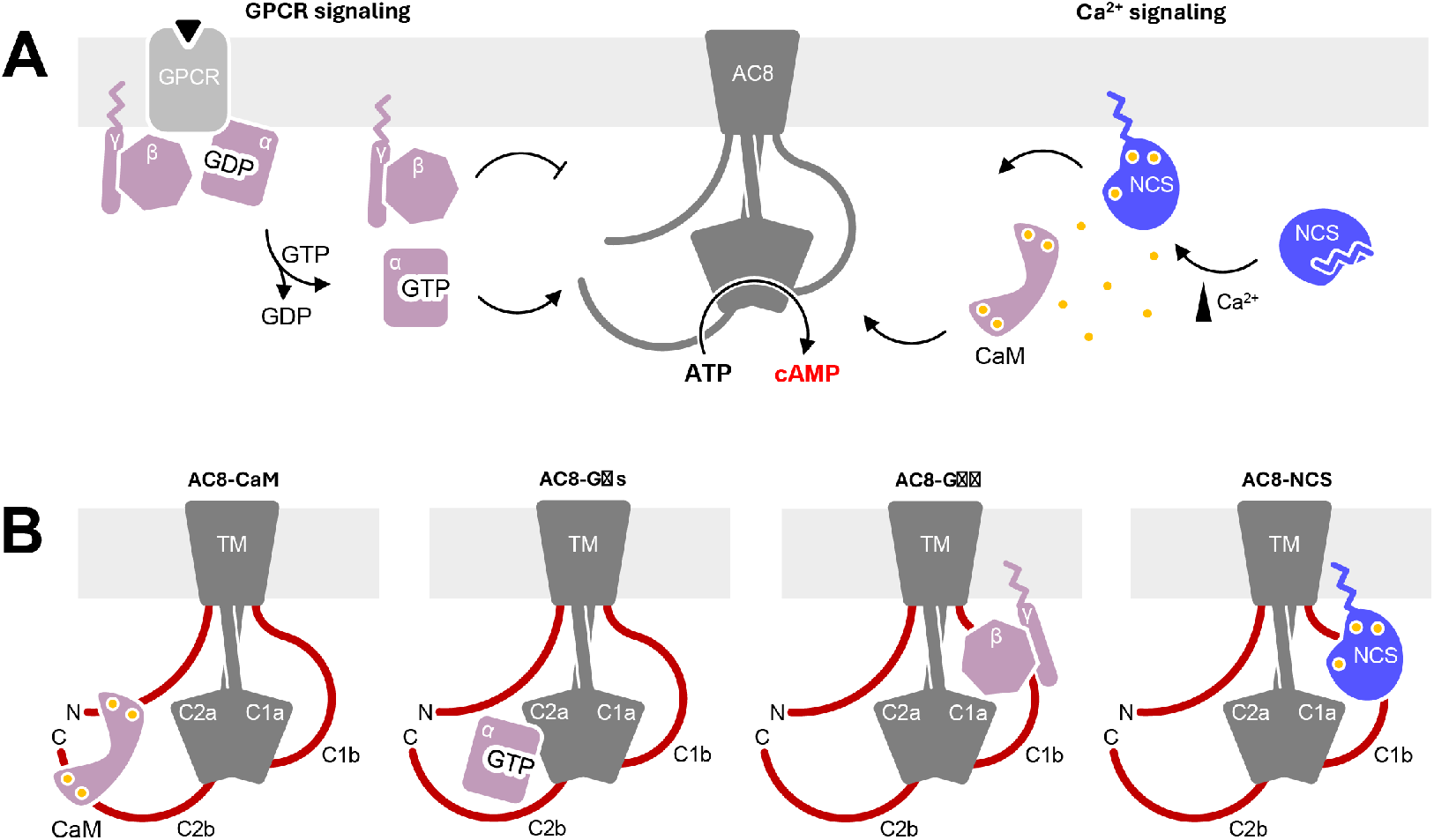
Schematic depiction of neuronal calcium sensor–mediated regulation of AC8. **N**euronal calcium sensor proteins interact with flexible N-terminal, C-terminal, and C1b domains, regulatory regions of AC8. Ca^2+^-dependent AC8-NCS interactions highlight novel context-dependent regulation of AC8 activity, as well as cross-talk between GPCR and Ca^2+^ signaling pathways. **(B)** Schematic models depicting binding mode of AC8-interactors. AC8 binds CaM via its N-terminal and the C2b domains and engages with Gαs via C2a, which results stimulated catalytic activity. Gβγ interact with the C1b domain, contributing to inhibition of catalytic activity. NCS proteins also interact with C1b domain, as well as with N- and C-terminal domains.

In addition, NCS proteins may act as molecular scaffolds, either recruiting AC8 to specific membrane compartments or facilitating the assembly or disruption of larger signalling complexes. This spatial regulation would allow for a highly restricted signaling response that is not easily captured in simplified *in vitro* assays [36]. AC8 has previously been found to be associated with lipid rafts, and disruption of AC8 glycosylation by mutagenesis influenced the enzymes localization to these microdomains[37]. Whereas some NCS proteins, such as neurocalcin, have been localized to lipid rafts[38], it has been anecdotally suggested that others, such as Vilip-1, do not specifically localize to rafts [39]. It is therefore tempting to suggest that interactions with myristoyl switch-containing NCS proteins may influence the distribution of AC8 in distinct lipid microdomains. As the flexible cytosolic regions in AC8 contain several putative phosphorylation sites, it is possible that myristoyl switch- and phosphorylation-dependent regulatory mechanisms control the association between NCS proteins and AC8 [40]. Defining the specific roles of NCS proteins in controlling lipid raft localization and trafficking of AC8 and other membrane ACs will require careful future experimentation.

Taken together, our findings support a model where NCS proteins act as context-dependent modulators of AC8. Within spatially organized signaling compartments, these sensors may tune AC8 activity by balancing inhibitory and stimulatory inputs, or by regulating the recruitment of additional signaling pathway components (**Figure 8**). In this framework, AC8 emerges as a multifunctional regulatory hub that integrates Ca^2+^/NCS signaling with GPCR-derived inputs in a spatially and temporally controlled manner.

## ACKNOWLEDGEMENTS

This work was supported by the grants to V.M.K. from Swiss National Science Foundation grants (184951 and 10003696) and from the Vontobel Stiftung (1669/2023).

## Competing interests

Authors declare that they have no conflict of interest.

## Author contributions

B.K., R.F., D.S., P.L., designed experiments, analysed data and prepared figures. B.K., R.F., D.S., P.L., A.K., M.R., I.K., J. S., V.M.K., performed experiments and analysed data. A.D.G., P.P., A.L., V.M.K. supervised the project. B.K., R.F., D.S., and V.M.K. wrote the manuscript with input from all co-authors.

## Data availability

The mass spectrometry proteomics data have been deposited to the ProteomeXchange Consortium (http://proteomecentral.proteomexchange.org) via the PRIDE partner repository [41] with the dataset identifiers PXD077139 for the tryptic digest, PXD056924 for the LiP-MS data and PXD075106 for the XL-MS data.

The AlphaLink2 models generated in this study based on experimental crosslinks have been deposited as integrative/hybrid models in PDB-IHM [42].

## Notes

### Competing Interest Statement

The authors have declared no competing interest.

### Summary of Updates

Illustrations / sketches in Figure 1A and Figure 8 revised and improved stylistically.

