## Supplementary Information for "Neuronal calcium sensors emerge as novel adenylyl cyclase 8 regulators"

### Material and Methods

***DNA constructs:*** The synthetic codon-optimized constructs of bovine AC8 and human neuronal calcium sensors (HPCAL1, HPCAL4, HPCA, VILIP, NCALD, NCS1) were purchased from Genewiz. All the neuronal calcium sensors were cloned in pACMV-based vectors with a C-terminal 10x-His tag, whereas bovine AC8 was cloned with a C-terminal YFP-Twin-Strep tag. The AC8 truncation constructs, AC8 -Nt (166-1253), AC8 -Ct (1-1172), AC8 -Nt -Ct (166-1172) were cloned into the pACMV-based vectors with C-terminal CFP or YFP tag using In-Fusion Snap Assembly (Takara).

**Protein expression and purification**

***Bovine AC8:*** Expression and purification of AC8 was performed using stably expressing HEK293S GnTi$^-$ monoclonal cell line as described in our previous study [1]. Briefly, HEK293S GnTI$^-$ cells expressing full-length bovine AC8 with C-terminal 3C-YFP-TwinStrep fusion tag were grown in FreeStyle 293 Expression medium. The expression was induced with tetracycline, and after 48-hour cell pellets were harvested and resuspended in 50 mM Tris-HCl pH 8.0, 150 mM NaCl, 10% glycerol supplemented with protease inhibitors (1 mM benzamidine, 1 µg/ml leupeptin, 1 µg/ml aprotinin, 1 µg/ml pepstatin, 1 µg/ml trypsin inhibitor and 1 mM phenylmethylsulfonyl fluoride (PMSF)). Cells were lysed using a Dounce homogeniser and solubilized in the 50 mM Tris-HCl pH 8.0, 150 mM NaCl, 10% Glycerol, 1% DDM and 0.02% cholesteryl hemisuccinate (CHS) for 1 hr at 4 °C. The detergent solubilised lysate was cleared by ultracentrifugation (Ti45 rotor, 186,000 × *g* for 40 min at 4 °C) and supernatant was subjected to GFP-nanobody affinity purification. The protein was eluted using cleavage by HRV 3 C protease (1:10 w/w) and subjected to size-exclusion chromatography (SEC) using a Superpose 6 Increase 10/300 GL column (GE Healthcare) equilibrated in 50 mM Tris-HCl pH 8.0, 150 mM NaCl, 0.02% GDN. The fractions corresponding to purified AC8 (elution volume, 13–16 mL) were collected, and flash frozen until use.

***HPCAL1 and other neuronal calcium sensors:*** Expression of HPCAL1 and other sensors were performed using transient transfection in HEK293F cells. All the NCS constructs contained C-terminal His-tag, with exception of HPCAL1-CFP construct with C-terminal strep tag. In brief, all the constructs were initially transformed into *E. coli* cells (MACH1) and the cells were grown on Luria–Bertani (LB) agar plates containing 100 µg/mL ampicillin. A single colony was picked to prepare an overnight culture either in LB or terrific broth (TB) containing 100 µg/mL ampicillin. The preculture was added to 800 mL LB or TB containing 100 µg/mL ampicillin and the cells were grown at 37°C for 16 hours. The cells were harvested at 4,500 rpm for 15 minutes (Sorvall BIOS 16) and stored at -80°C until further use. The plasmid was extracted using the NucleoBond Xtra Maxi kit (Macherey-Nagel) according to the manufacturer's instructions.

HEK293F cells grown to a density of 2 million cells/mL in 800 mL FreeStyle 293 Expression medium containing penicillin, streptomycin and amphotericin B were transfected with premixed 1.6 mg plasmid and 4.8 mg linear polyethyleneimine (40 kDa). The cells were harvested 3 days after transfection by centrifugation using Sorvall BIOS 16 centrifuge (1,000 x g, 20 minutes) and the cell pellets were frozen to -80°C and stored until further use.

All subsequent steps were carried out at 4°C. A cell pellet of 2.4 L transfected cells was resuspended in 50 mM Tris-HCl pH 8, 150 mM NaCl, 10% glycerol, containing 1 µg/mL pepstatin, 1 µg/mL leupeptin, 1 µg/mL aprotinin, 100 µg/mL trypsin inhibitor and 1 mM phenylmethylsulfonyl fluoride and lyzed using a dounce homogenizer. The lysate was solubilized with 1% DDM for 1 hour with slight agitation to solubilize membrane and membraneassociated proteins. The sample was centrifuged at 40,000 rpm for 40 minutes (Beckman Ti45 rotor) and the resulting supernatant was incubated with pre-equilibrated Ni-NTA beads for 45 minutes with slight agitation. The bead-sample mixture was loaded onto a 50 mL gravity column, the beads were washed with 20 column volumes 50 mM Tris-HCl pH 8, 150 mM NaCl, 5% glycerol, 0.02% GDN, 25 mM imidazole. In the second wash-step, the beads were washed with 20 column volumes 50 mM Tris-HCl pH 8, 150 mM NaCl, 5% glycerol, 0.02% GDN, 50 mM imidazole. The protein was eluted with 50 mM Tris-HCl pH 8, 150 mM NaCl, 5% glycerol, 0.02% GDN, 250 mM imidazole and the protein concentration of the eluted fractions was determined. The protein-containing fractions were concentrated and further purified by SEC using Superdex 200 Increase 10/300 GL column. The separated protein was further concentrated and flash frozen in liquid nitrogen after addition of 10% glycerol.

***Heteromeric G-proteins:*** The bovine Gαs subunit and Gβγ were expressed in Hi5 insect cells using the Bac-to-Bac baculovirus system, essentially as described previously[1]. Hi5 cells were grown in suspension to a density of $1.5 \times 10^6$ cells $mL^{-1}$ and infected with 1–2% P2 virus. Cells were harvested 72 h after infection. Cell pellets were resuspended, lysed and solubilized in 50 mM Tris-HCl pH 8.0, 150 mM NaCl, 1% DDM for 1 h. After clarification by ultracentrifugation (186,000 × g, 40 min, 4 °C), the supernatant was incubated with Ni-NTA resin for 30-60 min. The resin was washed first with 50 mM Tris-HCl pH 8.0, 150 mM NaCl, 0.02% DDM, and 20 mM imidazole, and then with 50 mM Tris-HCl pH 8.0, 150 mM NaCl, 0.02% GDN, 5% glycerol, and 40 mM imidazole. G-proteins were eluted in 50 mM Tris-HCl pH 8.0, 150 mM NaCl, 0.02% GDN, 5% glycerol, and 250 mM imidazole, concentrated using a 30-kDa cutoff concentrator, and further purified by SEC on a Superose 6 Increase 10/300 GL column equilibrated in 50 mM Tris-HCl pH 8.0, 150 mM NaCl, and 0.02% GDN. Peak fractions were concentrated, snap-frozen in liquid nitrogen, and stored at −80 °C.

**LC-MS/MS analysis**

***Protein digestion:*** Purified bovine AC8 was digested in triplicates. 3x20 µg of protein were diluted in 5% sodium deoxycholate and disulfide bonds were reduced with 5 mM tris(2-carboxyethyl) phosphine hydrochloride at 37°C, 800 rpm for 40 min. The free cysteines were alkylated by incubation with 40 mM iodoacetamide (room temperature, 800 rpm, in the dark, 30 min). The samples were diluted with 100 mM ammonium bicarbonate to a final sodium deoxycholate concentration of 1%. 0.5 µg of trypsin (Promega, V5113) and 0.5 µg of Lys-C (Fujifilm Biosciences, 129-02541) were added and the samples were incubated at 37°C, 800 rpm overnight. The next day, the digestion was stopped by adding formic acid to a final concentration of 2%. Precipitated sodium deoxycholate was removed by centrifugation. The obtained peptides were desalted and dried in a vacuum centrifuge. Prior to measurement, the samples were resuspended in 0.1% formic acid and transferred to HPLC vials.

***Data acquisition:*** The samples were measured on Orbitrap Eclipse Tribrid mass spectrometer (Thermo Scientific), equipped with a nanoelectrospray source and an Easy-nLC 1200 nanoflow LC system (Thermo Scientific). Peptides were separated on a 40 cm x 0.75 µm inner diameter column packed in-house with 1.9 um $C_{18}$ beads (Dr. Maisch Reprosil-Pur 120) using a linear gradient from 5-35% B (A: 0.1% FA, B: 95% ACN, 0.1% FA) over 30 min at 300 nL/min. The column was heated to 50 °C. The data was acquired with a data dependent acquisition (DDA) method. For MS1 spectra, Orbitrap resolution was set to 120,000, the scan range was between 375 and 1500 m/z. The RF lens voltage was set to 30%. Higher energy collisional dissociation (HCD) was performed on

5 precursors of charge states between +2 and +7 per cycle, with a normalized collision energy of 30% and an Orbitrap resolution of 30,000. The dynamic exclusion time was set to 60 s.

***Data analysis:*** The raw files were searched with SpectroMine v.4.4 (Biognosys), against a FASTA file containing the human proteome and the bovine AC8 sequence. An additional contaminants FASTA file based on the MaxQuant [2] contaminant database was included in the search. Normalization was switched off and single hit identifications were excluded. The list of detected proteins and their quantities were exported and further analyzed in R.

**HPCAL1 binding experiments**

Analysis of AC8 and NCS protein interactions was performed using SEC. For this, a total of 100 µg (0.71 µM) of purified AC8 with or without $Ca^{2+}$/NCS protein of interest (400 µg, ~17-18 µM) was injected on Superose 6 increase column pre-equilibrated in 25 mM Tris-HCl pH 8.0, 150 mM NaCl, 0.02% GDN, in the presence or in the absence of 1 mM $CaCl_2$. The fractions corresponding to the elution volume of AC8 (12-15.5 ml) and HPCAL1 (18-19.5 ml) were collected and analysed using SDS-PAGE. Interactions between AC8 and NCS proteins were inferred from their co-elution in the same SEC peaks.

AC8-HPCAL1 protein interaction analysis using fluorescence-detection size exclusion chromatography (FSEC) was performed in similar manner as described for AC8-CaM in our previous study [1]. In brief, the purified HPCAL1-CFP was used as a fluorescent probe and binding of AC8 with HPCAL1 was detected based on the appearance of CFP fluorescence at the position of the eluted AC8 peak. To estimate binding affinity, 14 ug of purified CFP-tagged HPCAL1 was mixed with various amounts of tag-free AC8 (at a range of concentrations, from 1.5 to 150 µg/ml) in a final volume of 100 µl. The mixtures were incubated on ice for 15 min, in the presence of $CaCl_2$ (1 mM) or EGTA (1 mM). After incubation, each sample was injected onto the Agilent SEC5-300 column (flow rate 0.3 ml/min, excitation 430 nm, emission 478 nm) pre-equilibrated in 25 mM Tris HCl, pH 8.0, 150 mM NaCl, 0.02% GDN with either 1 mM $CaCl_2$ or 1 mM EGTA. The total peak area (elution volume, 1.2-1.62 ml) and peak intensity (at 1.47 ml) were quantified for each experiment. The binding affinity of AC8 and HPCAL1 was evaluated using the one-site saturation binding model in GraphPad Prism 8.

**cAMP accumulation assay**

The cAMP accumulation assay was performed using an adapted protocol published by Alvarez and Daniels [3]. Briefly, purified AC8 at 10-15 nM and the AC8-NCS complex at 100 nM-100 µM NCS were prepared in 50 mM Tris-HCl pH 7.5, 150 mM NaCl, 0.02% GDN. In single-point comparative assays, G proteins and calmodulin were added at a 1.5-fold molar excess, whereas HPCAL1 was added at a 4-fold molar excess, relative to AC8 (10 nM). The reaction was started by mixing 100 µL AC8 or AC8-HPCAL1 with 100 µL of the 2X reaction mixture (4 mM $MgCl_2$, 2 mM $CaCl_2$, 10 mM $MgCl_2$, 0.2 mM ATP and 20 nM $^3$H-ATP (Perkin Elmer)). The mixture was incubated at 30°C for 30 min and quenched by addition of 20 µL 2.2 M HCl, followed by 4 min incubation at 95°C. The reaction mixture was cooled on ice and loaded onto gravity columns containing 1.3 g of aluminium oxide. The produced $^3$H-cAMP was eluted with 4 mL of 0.2 M ammonium acetate into scintillation vials, 12 mL scintillation liquid (LabLogic) was added into each vial and the amount of product was analyzed using Packard 2250 CA Tri-Carb liquid scintillation counter.

**Single particle cryo-EM**

The purified monomeric AC8 (~ca. 6 mg/mL) was incubated with 1 mM ATPαS, 0.1mM GTPγS, 1 mM CaCl2, 2 mM MgCl2, and 5 mM MnCl2, as well as a 2.5-fold molar excess of HPCAL1 and/or 1.2 molar excess of Gαs. Quantifoil 1.2/1.3 200-mesh grids were glow discharged in a PELCO easiGlow (Ted Pella) glow discharge system for 25 s at 30 mA. A 3.5 μL aliquot of the sample was deposited on the carbon side of the grid and blotted for 3 s (blot force 20) using a Vitrobot Mark IV (Thermo Fisher Scientific) with 100% humidity at 4°C, after which the grid was plunge-frozen in liquid ethane and stored in liquid nitrogen.

Initially, a small dataset of approximately 3000 movies was collected for AC8 alone, AC8-HPCAL1, AC8-Gαs and AC8-Gαs-HPCAL1. All movies were collected in super-resolution mode (EPU software) on a 300 kV Titan Krios electron microscope (Thermo Fisher Scientific) equipped with a Gatan K3 direct electron detector and a Gatan Quantum-LS GIF at ScopeM (ETH Zurich). The movies were acquired at a defocus range of -0.5 to -3 μm. The dose per movie was 55 $e^{-}$/ $Å^2$ and all movies were two-fold binned resulting in a pixel size of 0.66 Å. Additional 3 datasets for AC8-HPCAL1 complex (8974, 13588, and 11360 movies with total dose of 50 $e^{-}$/ Å2, 67 $e^{-}$/ Å2, and 60 $e^{-}$/ Å2, respectively), were collected. An additional dataset of 6363 movies with a total dose of 50 $e^{-}$/ Å2 was acquired for AC8-Gαs-HPCAL1 complex.

Data processing was performed with Relion (v. 3.1.3). All movie stacks were motion corrected using MotionCorr (v. 1.1.0). The micrographs were corrected by CTF (GCTF). 1211 particles were picked manually to generate a template for autopicking. A fraction of 42,689 particles showed a clear density for HPCAL1, these were used for model generation and 3D classification. After several rounds of 3D classification, a 3D reconstruction map at 21 Å could be obtained.

**Limited proteolysis-coupled mass spectrometry**

Membrane preparations were produced as described elsewhere [1, 4]. The membrane suspensions of AC8-expressing HEK293 GnTI$^{-}$ cells were diluted with 100 mM HEPES-KOH pH 7.4, 150 mM KCl, 1 mM $MgCl_2$, 1 mM CaCl2 and 36 aliquots (50 μL each at 2 μg/μL protein) were prepared. Purified HPCAL1 was diluted with 100 mM HEPES-KOH pH 7.4, 150 mM KCl, 1 mM $MgCl_2$, 1 mM $CaCl_2$, 0.02% GDN. All experiments were conducted in quadruplicates. Serial dilutions of HPCAL1 were spiked into the samples in increasing concentrations (0 μg, 0.01 μg, 0.1 μg, 0.5 μg, 1 μg, 2 μg, 3 μg) and they were incubated for 10 min at 25°C. Limited proteolysis was performed by adding 1 μg of proteinase K, or the same volume of water for tryptic controls (only for the highest and lowest HPCAL1 concentrations), to the samples and incubating them at 25°C for 5 minutes. The samples were then incubated for 5 min at 99°C, followed by cooling to 4°C and the addition of an equal volume of 10% sodium deoxycholate to quench the reaction.

All samples were transferred to a 96-well plate and disulfide bonds were reduced by adding 5 mM tris(2-carboxyethyl)phosphine hydrochloride and incubating at 37°C for 40 minutes with slight agitation. Free cysteines were alkylated by adding 40 mM iodoacetamide, followed by an incubation at room temperature in the dark with slight agitation. The samples were diluted with 100 mM ammonium bicarbonate to achieve a sodium deoxycholate concentration of 1%. 1 μg of trypsin and 1 μg of Lys-C were added to the samples, followed by an incubation at 37°C overnight with slight agitation. The next day, the digestion was stopped by adding formic acid to a final concentration of 2%, which led to sodium deoxycholate precipitation. The precipitated sodium deoxycholate was removed by filtration (96-well Corning® 2 µm PVDF plate). The resulting peptides were desalted using a 96-well MacroSpin C18 plate (The Nest Group). Peptides were eluted with 80% acetonitrile, 0.1% formic acid and dried in a vacuum centrifuge. The 96-well plate containing the dried peptides was stored at -20°C until further use.

Prior to LC-MS/MS analysis, the peptides were resuspended in 40 µL 5% acetonitrile, 0.1% formic acid with addition of iRT peptides (Biognosys). An aliquot of each sample of each condition was pooled with the other replicates to prepare samples for spectral library generation. 1 µL of each sample was injected an Easy nLC 1200 nanoflow liquid chromatography system equipped with a 40 cm in-house packed column (1.9 µm beads (Dr. Maisch Reprosil-Pur 120), heated to 50°C. Peptides were separated with a linear gradient from 3-30% eluent B (eluent A: 0.1% formic acid, eluent B: 95% acetonitrile, 0.1% formic acid) over 120 min at 300 nL/min. Library samples were acquired with data dependent acquisition (DDA), the other samples were acquired with data independent acquisition (DIA) on an Orbitrap Exploris 480 mass spectrometer (Thermo Scientific). Both, the DDA and the DIA method, included a lock mass for internal mass calibration (445.12003 m/z). For DDA measurements, the Orbitrap resolution for full scans (MS1) was set to 120,000 at a scan range between 350-1150 m/z. The RF lens was set to 40%, the normalized AGC target was set to 200%. The maximum injection time was set to 100 ms. Dynamic exclusion was set to 20 s and charge states 2-6 were included. For data dependent MS2 scans selected peptides were fragmented with higher energy collisional dissociation at a collision energy of 30%. Orbitrap resolution was set to 30,000, normalized AGC target was set to 200%, maximum injection time was 54 ms. For DIA measurements the Orbitrap resolution was set to 120,000, the scan range was 350-1150 m/z. RF lens was set to 50%, the normalized AGC target was set to 200%. The maximum injection time was 264 ms. The DIA method was set up with 41 variable width windows (1 m/z overlap). The fragmentation mode was higher energy collisional dissociation with a collision energy of 30% and an Orbitrap resolution of 30,000. The normalized AGC target was 2,000%, the maximum injection time - 66 ms.

The acquired raw data was searched using Spectronaut v. 15.4 (Biognosys). The tryptic control and LiP data were searched separately. A library was produced from searches of DDA data with a slight modification of the default parameters: minimum peptide length was set to 5 amino acids, single hits were excluded, and missing values were not imputed. For LiP-MS data, protease specificity was set to "semi-specific". Data was exported from Spectronaut and further analyzed in R. Quality control, data analysis and interpretation was conducted using the R packages protti[5], tidyverse[6], data.table[7]. The tryptic control data was checked for AC8 protein abundance changes and four-parameter dose response curves were modeled using the intensity of all detected AC8 peptides in the LiP samples with an implementation of the drc [8] R package within protti. Pearson correlation was calculated and peptides with a Pearson correlation r of >0.85 were considered significant. Peptides with only one changing concentration point were not considered significant.

**Cross-linking and sample preparation for mass spectrometry**

AC8 and the neuronal calcium sensors HPCAL1, NCALD and VILIP1 were purified in crosslinking buffer (20 mM HEPES pH 7.5, 150 mM NaCl, 10% Glycerol, 0.02% GDN). Other calcium sensors were buffer exchanged into the crosslinking buffer using a Zeba Spin desalting column, 7K MWCO (ThermoFisher Scientific). For each experimental condition, 25 µg each of AC8 and 25 µg of corresponding NCS were added. The proteins were incubated on ice for 20 minutes in the cross-linking buffer supplemented with final concentrations of 0.5 mM

$CaCl_2$, 1 mM $MgCl_2$ and 2.5 mM $MnCl_2$. All the conditions were filled up to 200 ul total, corresponding to 0.25 mg/ml concentration. Cross-linking was then performed as previously described [9].

To cross-link primary amines, a 1:1 mixture of DSS-$d_0$ and DSS-$d_{12}$ (Creative Molecules; at a 25 mM stock concentration in dimethylformamide) was added to the sample, bringing the final concentration to 1 mM. The sample was then incubated for 30 min at 25°C with mild shaking (750 rpm on the thermomixer). To quench the reaction, 1 M ammonium bicarbonate (ABC) was added to achieve a final concentration of 50 mM. The mixture was incubated for a further 20 minutes at 37 °C with gentle shaking at 750 rpm. The samples were then dried by evaporation. Dried samples were resuspended in 8 M urea to a final concentration of 1 mg/ml. The disulfide bonds were reduced by adding tris(2-carboxyethyl) phosphine to a final concentration of 2.5 mM. The samples were first incubated for 30 minutes at 37°C, then cooled to room temperature before carbamidomethylation with iodoacetamide, which was added to a final concentration of 5 mM. After a 30-minute incubation in the dark at room temperature, the samples were diluted with 150 mM ABC to a final urea concentration of 5.5 M. Endopeptidase Lys-C (Wako) was added at an enzyme-to-substrate ratio of 1:100 and the samples were incubated for two hours at 37 °C with mild shaking (750 rpm). The samples were then further diluted to 1 M urea using 50 mM ABC before trypsin was added at an enzyme-to-substrate ratio of 1:50. The samples were digested overnight at 37 °C with mild shaking (750 rpm). Digestion was stopped the following day by adding 100% formic acid to achieve a final concentration of 2%. The samples were desalted using solid-phase extraction (Sep-Pak tC18 cartridges, Waters) and then fractionated using peptide-level SEC (Superdex 30 Increase column, 300 x 3.2 mm; GE Healthcare). The mobile phase consisted of water/acetonitrile/trifluoroacetic acid (70:30:0.1, v/v/v) at a flow rate of 50 µl/min. Four 100-µl fractions, corresponding to an elution volume of 1.0–1.4 ml, were collected from each sample. The fractions were dried by evaporation in a vacuum centrifuge.

**Liquid chromatography-tandem mass spectrometry (LC-MS/MS)**

Chromatographic separation was done on an Easy nLC-1200 HPLC system coupled to an Orbitrap Fusion Lumos mass spectrometer via a Nanospray Flex ion source (all manufactured by Thermo Fisher). SEC fraction were injected in duplicates and separated on an Acclaim PepMap RSLC C18 column (250 mm × 75 µm, 2-Å particle size, Thermo Fisher Scientific). The mobile phase consisted of (A) 0.15% formic acid in water/acetonitrile (98:2, v/v) and (B) 0.15% formic acid in acetonitrile/water (80:20, v/v). Peptides were eluted at 300 nL/min using a 60-minute linear gradient from 11% to 40% B.

Mass spectra were collected in data-dependent acquisition (DDA) mode with a 3-second cycle duration. Full MS scans were captured in the Orbitrap at 120,000 resolution, while MS/MS fragmentation was recorded at 30,000 resolution. Precursors (3+ to 7+ charge; m/z 350–1500) were fragmented by collision-induced dissociation (CID) in the linear ion trap (35% normalized collision energy). A 30-second dynamic exclusion was applied post-sequencing.

**Identification of cross-linked peptides**

Cross-linked peptides were identified using xQuest (version 2.1.5, https://gitlab.ethz.ch/leitner_lab/xquest_xprophet [10]). A database including the proteins of interest and the most abundant contaminants (a total list of 10 proteins) was used to search the data. A decoy database containing the reversed sequences was also included. The search parameters comprised trypsin as the enzyme; a maximum of two missed cleavages; carbamidomethylation of Cys as the fixed modification; oxidation of Met as the variable modification; an MS1 error tolerance of ±15 ppm; and an MS2 error tolerance of ±15 ppm. A filtering step was then applied to the cross-linked peptide candidates, with the following parameters: a mass tolerance window of −5 to 5 ppm, a delta score of <0.9, and a minimum of five matched fragment ions per peptide. The resulting datasets correspond to a false discovery rate of less than 1% at the unique peptide-pair level. The obtained list of crosslinks was further filtered and prepared for visualization using xiView [11].

**NMR Spectroscopy**

***M9 medium for U-[$^{15}N$] isotope labelling :*** To prepare 1 L of medium, 7.5 g/L $Na_2HPO_4·2H_2O$, 0.5 g/L NaCl, 4 g/L D-glucose, 1 µg/mL thiamine, 1 µg/mL biotin, 1 mM $MgSO_4$, 0.33 mM $CaCl_2$, 1 mL of 1k trace elements, 100 µL of 10k trace elements, and 50 µg/mL kanamycin were added. For preparation of a stock solution of 1k trace elements, 0.1 mg/mL $H_3BO_3$, 0.84 mg/mL $ZnCl_2$, 0.1 mg/mL $CoCl_2·6H_2O$, 0.13 mg/mL $CuCl_2·2H_2O$, 8.33 mg/mL $FeCl_3·6H_2O$, and 50 mg/mL ethylenediaminetetraacetic acid (EDTA) were dissolved in 100 mL dd$H_2O$. For preparation of a stock solution of 10k trace elements, 33.7 mg/mL $CuSO_4·5H_2O$, 5 mg/mL $CoCl_2·6H_2O$, 18 mg/mL $MnSO_4·H_2O$, and 4.3 mg/mL $ZnSO_4·7H_2O$ were dissolved in 100 mL dd$H_2O$. Finally, 1 g/L $^{15}NH_4Cl$ was dissolved and the medium was sterile filtered through a 0.22 µm filter.

***Expression of U-[$^{15}N$] labelled peptides in E.coli :*** GB1-6xHis-bAC8-Nt1 (bovine AC8 residues 1–95), GB1-6xHis-bAC8-Nt2 (bovine AC8 residues 70–164), GB1-6xHis-bAC8-C1b1 (bovine AC8 residues 592–690), GB1-6xHis-bAC8-C1b2 (bovine AC8 residues 631–710), and GB1-6xHis-bAC8-Cter (bovine AC8 residues 1166–1253) plasmids were transformed into BL21 E. coli cells and grown on LB-agar plates containing 50 µg/mL kanamycin at 37 °C. A single colony was picked to prepare a preculture in 100 mL LB containing 50 µg/mL kanamycin and was grown for 4 hours at 30 °C with constant shaking at 180 rpm. The culture was centrifuged at 4000 rpm for 20 minutes and the supernatant was discarded. The pellet was resuspended in 100 mL of M9 medium enriched with 1 g/L $^{15}NH_4Cl$ and grown at 30 °C with constant shaking at 180 rpm. Once the culture reached $OD_{600}$ 1.0, it was re-centrifuged at 4000 rpm for 20 minutes, the supernatant was discarded, and the pellet was resuspended in 1 L of M9 medium enriched with 1 g/L $^{15}NH_4Cl$ and grown at 37 °C with constant shaking at 180 rpm. After the $OD_{600}$ reached 0.7–0.8, the incubation temperature was reduced to 20 °C. Expression was induced by adding isopropyl β-D-1-thiogalactopyranoside (IPTG) to a final concentration of 1 mM. The cells were harvested the following day at 4500 rpm for 15 minutes at 4 °C and stored at −20 °C before use.

***Peptide purification**:* Bacterial cell pellets were resuspended in 50 mM Tris-HCl pH 8.0, 300 mM NaCl, 10% glycerol, and 1 mM phenylmethylsulfonyl fluoride (PMSF), and then sonicated for 1 min (0.5 cycle ON / 0.5 cycle OFF). The lysate was cleared by centrifugation at 12,000 rpm for 30 min and incubated with Ni-NTA resin for 1 hour at 4 °C with slight agitation. The resin was washed with 12 column volumes of wash buffer containing 50 mM Tris pH 8.0, 300 mM NaCl, 10% glycerol, and 10 mM imidazole. The peptides were eluted in 50 mM Tris pH 8.0, 300 mM NaCl, 10% glycerol, and 250 mM imidazole. 1 mM dithiothreitol (DTT) was added to the eluate. The peptide-containing fractions were pooled, concentrated with 10 kDa molecular weight cut-off concentrators, and further purified using size-exclusion chromatography (Superdex 200 Increase 10/300 GL) in 50 mM HEPES pH 7.0, 150 mM NaCl, 0.5 mM $CaCl_2$, 1 mM $MgCl_2$, and 0.02% glyco-diosgenin (GDN).

***Data collection and analysis****:* All spectra were obtained on a 500 MHz spectrometer (Bruker) equipped with a $^{1}H$-$^{13}C$/$^{15}N$/D TCI Cryoprobe (Bruker) with z-gradients in 50 mM HEPES pH 7.0, 150 mM NaCl, 0.5 mM $CaCl_2$, 1 mM $MgCl_2$, and 0.02% GDN. All measurements were performed at 283 K in 3 mm sample tubes. All NMR samples were supplemented with 10% $^{2}H_2O$ for the deuterium lock. 150 µL of ~50 µM fusion peptide was used to record $^{1}H$–$^{15}N$ spectra of each fusion peptide alone. For interaction experiments, purified HPCAL1 was added to a final molar ratio of 1:1.26 (fusion peptide:HPCAL1). XL-ALSOFAST-[$^{13}C$,$^{1}H$]-HMQC spectra were recorded with TD (direct) = 2048 and TD (indirect) = 256, SI (direct) = 2048 and SI (indirect) = 1024, NS = 128, O1 (indirect) = 120.000 ppm, and D1 = 0.2 s to observe potential interactions.

**Cell culture for light microscopy experiments**

HEK293F cells were maintained in Dulbecco's Modified Eagle Medium (DMEM) with 10% fetal calf serum (FCS) and 1% penicillin, streptomycin (PenStrep) at 37°C, at 5% CO2. For calcium-stimulation, HEK293F cells were seeded in ibidi µ-slide microscopy chambers, precoated with poly-L-lysine. The cells were seeded at 40,000 cells/well in DMEM with 10% FCS, 1% PenStrep. The cells were transfected with 1 µg of the G-GECO $Ca^{2+}$-sensor construct at a 1:2 ratio of DNA to branched polyethyleneimine (w/w). For FRET analyses, the CFP and YFP constructs were co-transfected at a ratio of 2:1 (CFP: YFP). The cells were incubated at 37°C, 5% CO2, for 48 h.

**Light microscopy protein expression analysis**

Each well was washed with phosphate buffered saline (pH 7.4) and 200 µL of FluoroBright DMEM supplemented with 1% FCS and PenStrep. The cells were imaged on a Leica Stellaris 5 microscope, using LAS X software (4.2.1.23819 - build 23180), using an HC PL APO 63X/1.4 oil CS2 objective and 405 nm and 488 nm laser lines. 10 images were collected per condition with the same imaging parameters.

**Time-resolved FRET analyses**

The cells were washed with phosphate buffered saline (pH 7.4) and incubated with 100 µL phosphate buffered saline (pH 7.4) for 10 min at 37°C, 5% CO2. The sample was placed in the Leica Stellaris 5 microscope, equipped with two HyD detectors. The time-resolved FRET series were acquired using the FRET SE module (using LAS X software; 4.2.1.23819 - build 23180) according to the user manual using an HC PL APO 63X/1.4 oil CS2 objective and 405 nm and 488 nm laser lines to excite CFP (donor) and YFP (acceptor) fluorophores, respectively. For the ionomycin stimulation experiment, an image was acquired every 1.2 seconds for 6 min. The calcium increase was stimulated after the first micrograph using a final concentration of 3 µM ionomycin with 10 mM CaCl2.

The HPCAL1 redistribution was measured from the donor-only channel and was analyzed with Fiji (ImageJ) by measuring the intensities at the plasma membrane and the cytoplasm over time in separate regions of interest. The intensities were plotted over time and normalized between 0 and 100%. The data was analyzed using LAS X software by measuring the FRET efficiency at the plasma membrane of each cell according to [12]:

FRET efficiency = (FRET channel-Donor channel * $\beta$-Acceptor channel *($\gamma$-$\alpha$ * $\beta$)) / (Acceptor channel *(1- $\beta$ * $\delta$))

α, β, γ, δ are calibration factors generated by the acceptor and donor only references. The FRET efficiency values were plotted over time using GraphPad Prism (v. 9.0.0) with one-phase and two-phase association curve fits.

**Structural Modelling using AlphaFold3**

AC8-NCS complexes were predicted using AlphaFold 3.0.1 [13]. Protein sequences were obtained from UniProt[14] and ligand structures were defined using SMILES strings . The models were run on Euler scientific computing cluster (ETH Zurich). All protein complexes were predicted in the presence of 3 calcium ions explicitly defined as ligands. A total of 100 predicted models were generated using a combination of 20 random seeds and five recycling iterations per seed. The models were then ranked by ipTM and pTM scores.

**Structural Modelling with AlphaLink2**

Experimental distance restraints were derived from crosslinking experiments described above, only crosslinks between two proteins, were used for predictions. Predictions were performed using the AlphaLink2 framework, which extends the Uni-Fold-Multimer architecture to integrate sparse experimental data directly into the pair representation [15-17]. The models were run on Euler scientific computing cluster (ETH Zurich). Protein sequences were obtained from UniProt[14] and the AlphaLink-Multimer_SDA_v3 model weights were used to predict all complexes with 20 random seeds and five recycling iterations, producing 100 predictions in total. All crosslinks were applied at a 20% FDR for the predictions. The PDB template database was set up to May 2024. All the other settings are as described on the GitHub page (https://github.com/Rappsilber-Laboratory/AlphaLink2). The resulting models were all given a rank based on overall confidence, ipTM, pTM, and manual inspection of the model. The top five hits were further scored by PAE and pLDDT in predicted binding sites. Finally, the cross-links were mapped back to the models to see if the model matches the distances of the cross-links that were determined experimentally.

### Supplementary Figures and Tables

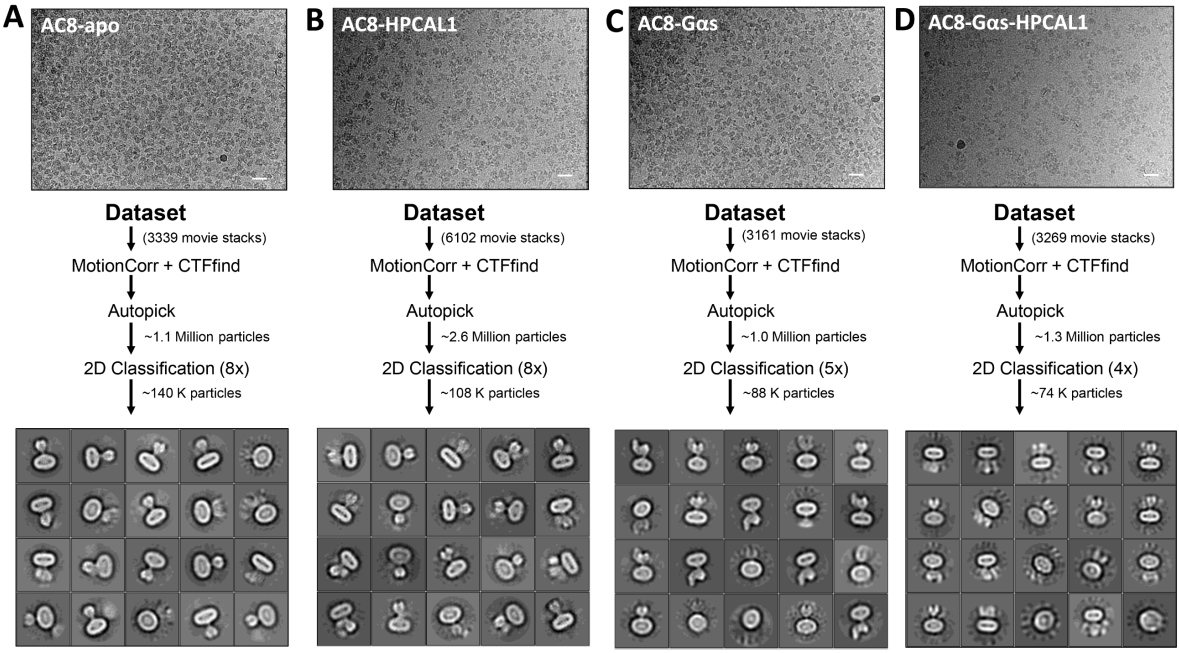


**Supplementary Fig. S1**. Cryo-EM analysis of small datasets of AC8 or AC8-Gαs complexes in presence and absence of HPCAL1. The representative micrographs and 2D class averages of **(A)** AC8 alone, **(B)** AC8-HPCAL1, **(C)** AC8-Gαs and **(D)** AC8-Gαs-HPCAL1. The scale bar for micrographs is 20 nm.

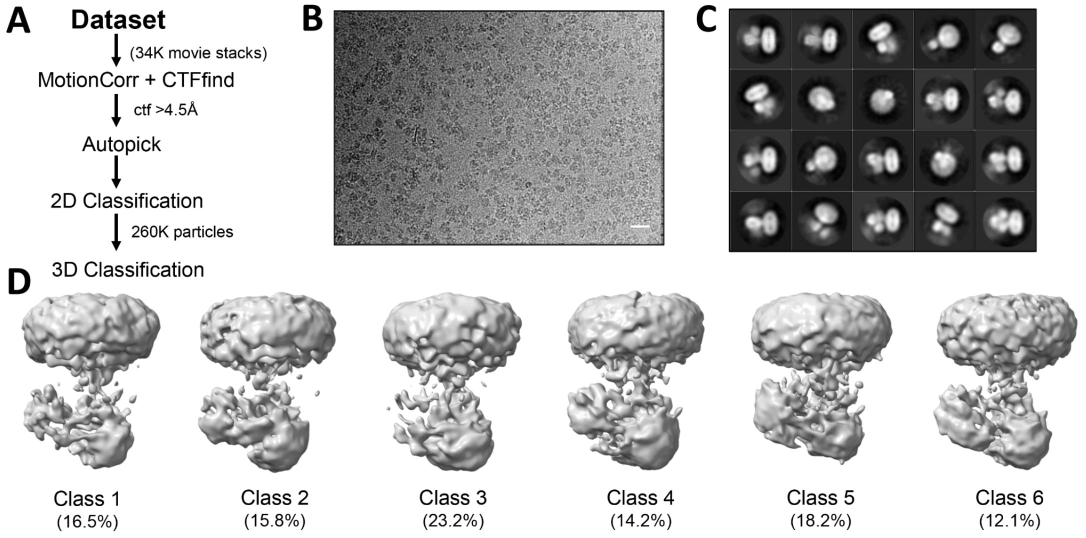


**Supplementary Fig. S2**. Cryo-EM analysis of AC8-HPCAL1 (large datasets). (**A**) Schematic representation of the cryo-EM data processing pipeline. **(B)** Representative micrographs and **(C)** 2D class averages. **(D)** Representative 3D classification jobs depicting low resolution 3D maps for each class.

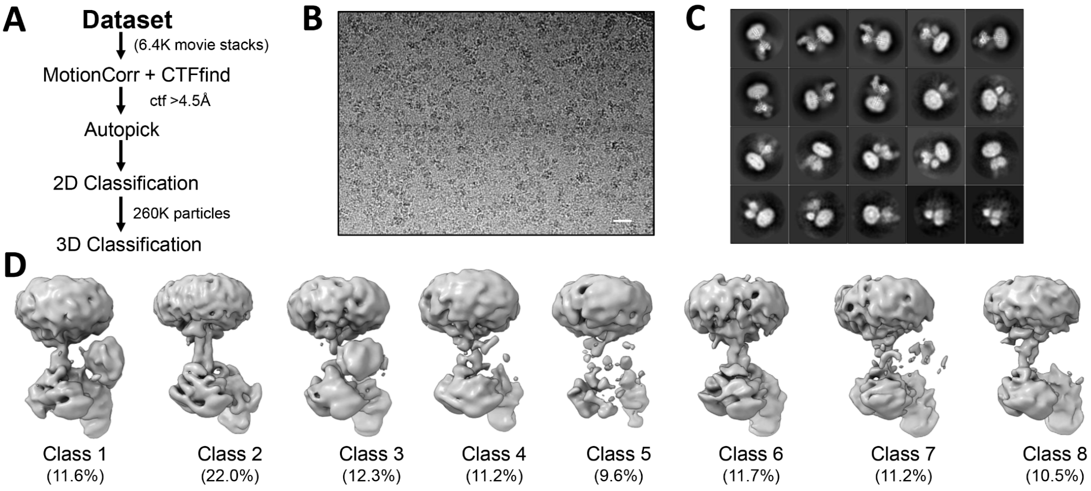


**Supplementary Fig. S3**. Cryo-EM analysis of AC8-Gαs-HPCAL1 (large datasets). **(A)** Schematic representation of the cryo-EM data processing pipeline. **(B)** Representative micrographs and **(C)** 2D class averages. **(D)** Representative 3D classification jobs depicting 3D maps for each class.

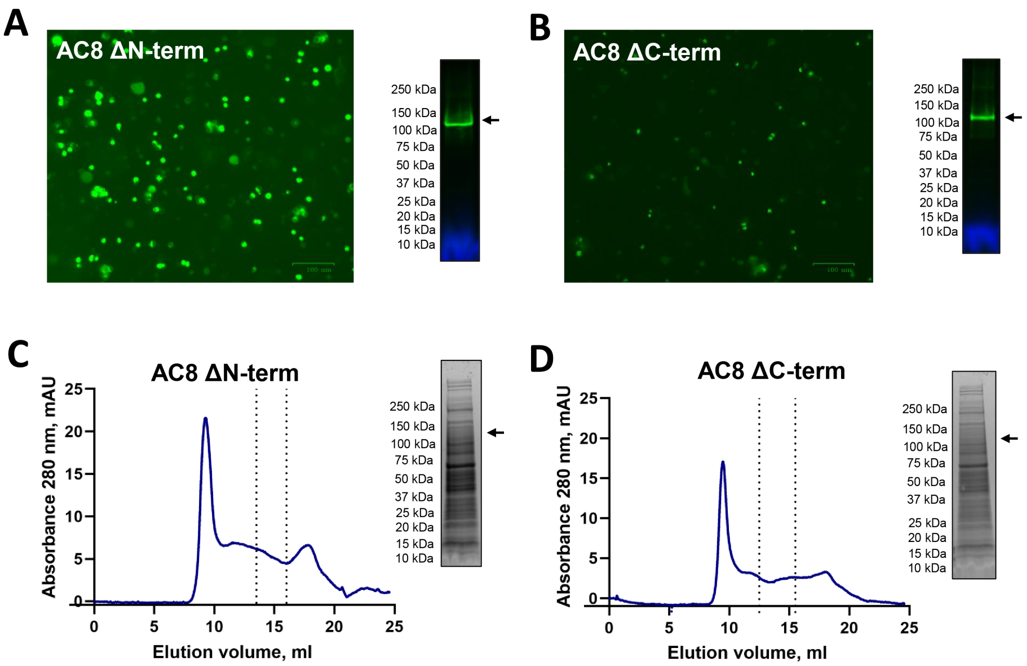


**Supplementary Fig. S4.** Expression and purification attempt of AC8 deletion constructs. **(A–B)** Fluorescence microscopy images showing expression of AC8 constructs lacking either the N-terminus (AC8ΔN-term) or the C-terminus (AC8ΔC-term), each fused to a C-terminal YFP tag. Insets on the right of each panel display in-gel fluorescence of detergent-solubilized membrane fractions, confirming expression of intact YFP-tagged AC8 constructs. **(C–D)** Size-exclusion chromatography (SEC) chromatograms and corresponding SDS–PAGE analysis of eluted fractions for AC8ΔN-term and AC8ΔC-term constructs. No clear protein peak or recoverable fraction corresponding to AC8 was observed, indicating instability and/or loss of the protein during purification.

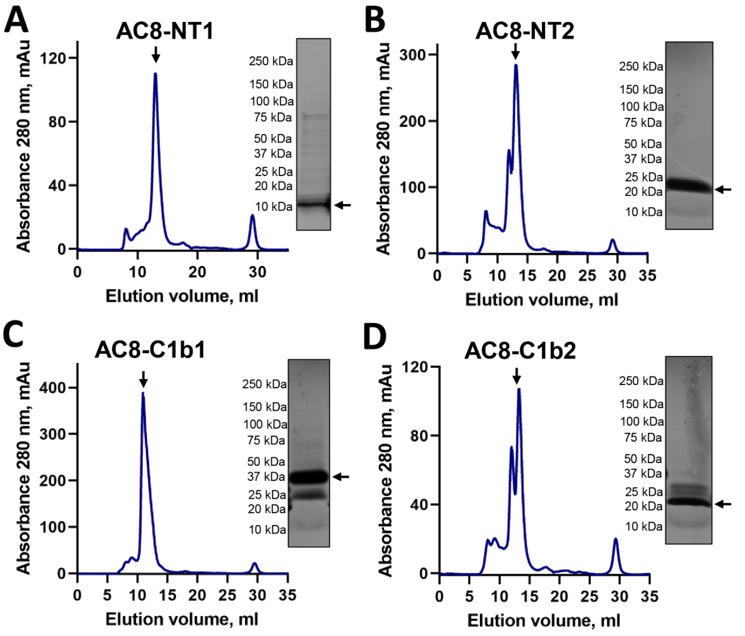


**Supplementary Fig. S5**. Size-exclusion chromatography profiles and corresponding SDS–PAGE analyses of purified (A-B) AC8 N-terminal fragments (AC8-NT1 and AC8-NT2) and (C-D) C1b fragments (AC8-C1b1 and AC8-C1b2) employed for NMR experiments.

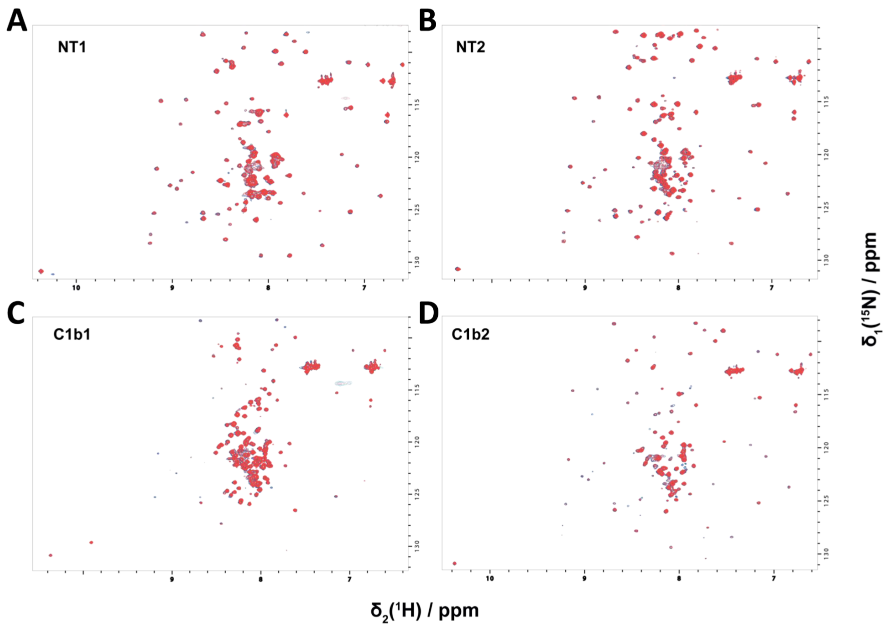


**Supplementary Fig. S6**. NMR analysis shows no detectable direct interaction between HPCAL1 and isolated N-terminal and C1b regions of AC8. Overlay of two-dimensional $^{1}H$–$^{15}N$ HSQC spectra of N-terminal domain NT1 fragments (A), NT2 (B) and C1b domain fragments C1b1 (C) and C1b2 (D) recorded in the absence (blue) and presence (red) of increasing concentrations of HPCAL1. No significant chemical shift perturbations or line broadening were observed upon addition of HPCAL1, indicating the absence of stable or high-affinity interactions with isolated AC8 N-terminal or C1b regions under these conditions.

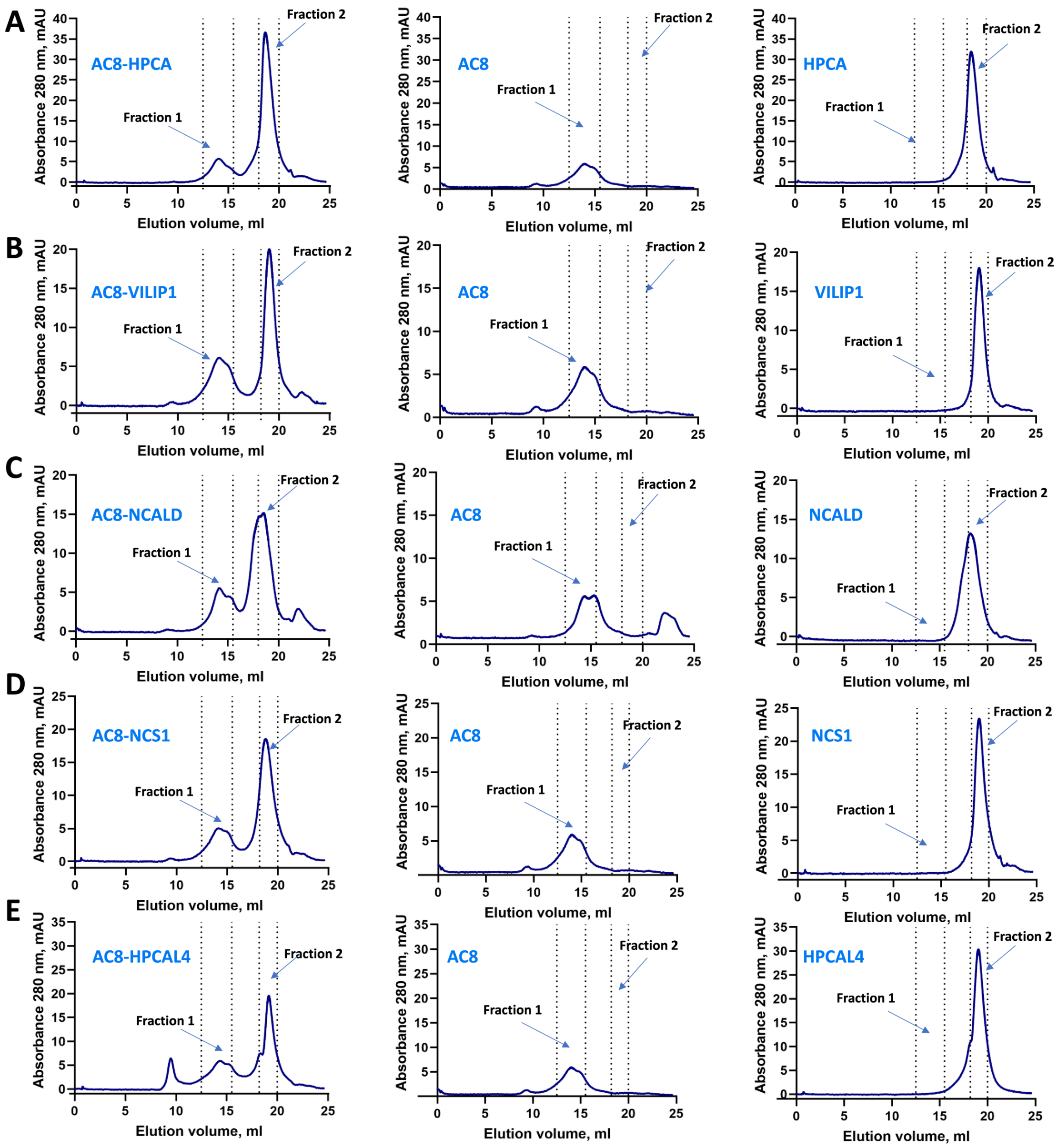


**Supplementary Fig. S7**. Representative profiles in size-exclusion chromatography (SEC) -based AC8 binding experiments for (A) HPCA (B) VILIP1 (C) NCALD (D) NCS1 (E) HPCAL4. Each experiment was performed by injecting purified AC8 alone, neuronal calcium sensors alone and pre-incubated mixtures of them as shown in each panel.

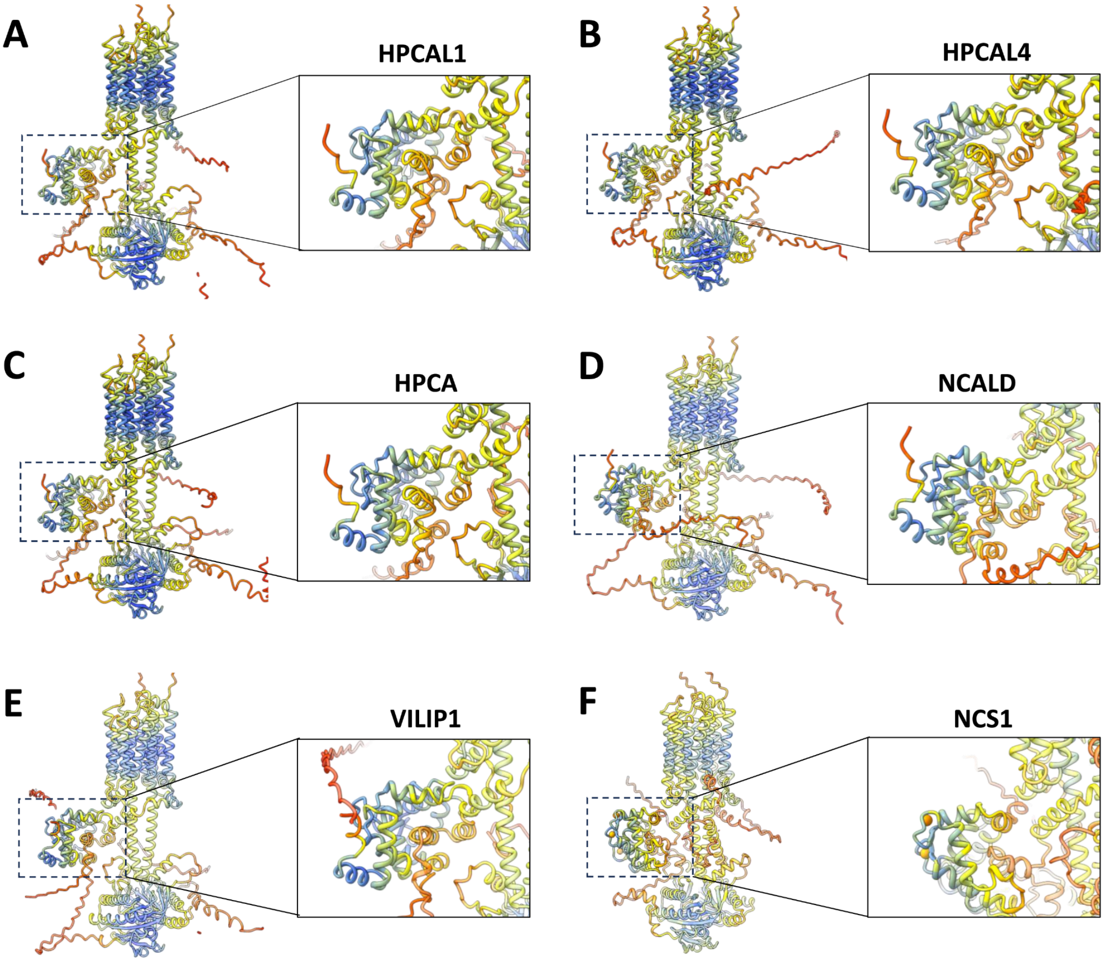


**Supplementary Fig S8** AlphaLink2 models of (A) HPCAL1 (B) HPCAL4 (C) HPCA (D) NCALD (E) VILIP1 and (F) NCS1. Model confidence is indicated by color according to predicted local distance difference test (pLDDT) scores: dark blue, very high confidence (pLDDT > 90); light blue, high confidence (70 < pLDDT ≤ 90); yellow, low confidence (50 < pLDDT ≤ 70); orange, very low confidence (pLDDT ≤ 50).

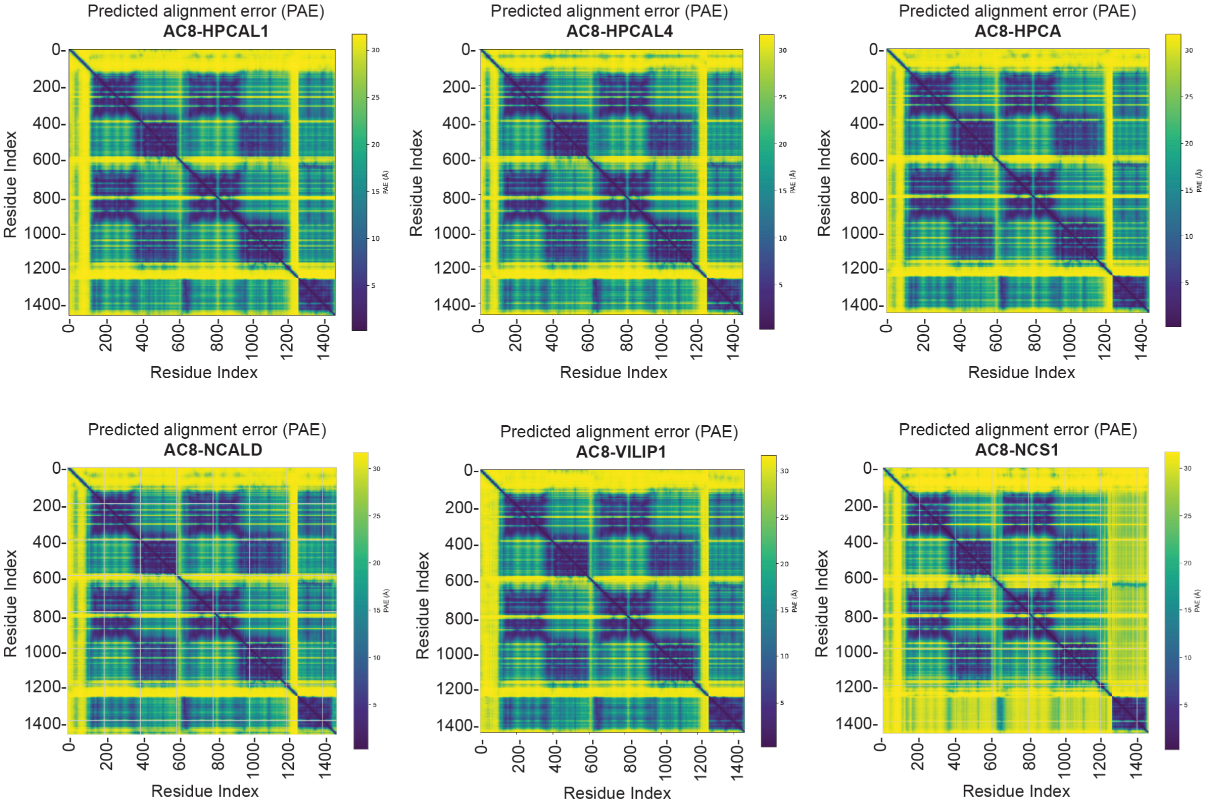


**Supplementary Fig S9.** Predicted alignment error (PAE) matrices for AlphaFold/AlphaLink models of AC8 in complex with HPCAL1, HPCAL4, HPCA, NCALD, VILIP1, and NCS1. Each heat map displays the expected positional error (in Å) between residue pairs, with darker colors indicating lower predicted error (higher confidence) and lighter colors indicating higher uncertainty. Axes correspond to residue indices spanning the full-length AC8 (1253 amino acids) – NCS complex. Diagonal blocks of low PAE reflect high confidence within individual protein domains, whereas off-diagonal regions report confidence in inter-domain and inter-protein orientations.

**Supplementary Table 1.** Comparison of AlphaFold and AlphaLink model confidence scores for AC8–neuronal calcium sensor complexes.

| | **AlphaFold 3** | | **AlphaLink2** | |
|---|---|---|---|---|
| | **pTM** | **ipTM** | **pTM** | **ipTM** |
| **HPCAL1** | 0.55 | 0.59 | 0.63 | 0.67 |
| **HPCAL4** | 0.61 | 0.55 | 0.64 | 0.68 |
| **HPCA** | 0.55 | 0.47 | 0.64 | 0.69 |
| **VILIP1** | 0.59 | 0.54 | 0.65 | 0.69 |
| **NCALD** | 0.58 | 0.51 | 0.65 | 0.67 |
| **NCS1** | 0.58 | 0.48 | - | - |
